# Low-grade inflammation in obesity causes low-effort food choice

**DOI:** 10.64898/2026.01.09.698429

**Authors:** Judith M. Scholing, Rinke Stienstra, Lisette Olsthoorn, Marijn S. Hendriksz, Catharina M. Mulders-Manders, Ruben van den Bosch, Esther Aarts

## Abstract

Inflammation relates to decreased effortful behaviour and altered effort-related responses in brain regions, such as the dorsomedial prefrontal cortex (dmPFC). Inflammation is prevalent in obesity, but its effects on effortful versus more convenient food choices are unknown. We investigated the role of low-grade inflammation in effortful food choice using functional MRI in a cross-sectional study (n=150 women, BMI>27 kg/m²) and a 12-week randomized controlled trial (n=59 women, BMI>30 kg/m², CRP>3 mg/L; colchicine 0.5 mg/d vs. placebo). Inflammation was related to less high-effort choices (OR=0.27, p=0.004) and higher effort-related dmPFC signal (β=0.23, p=0.025, Rpartial²=0.039), and mediated the association between BMI and dmPFC signal. Colchicine decreased systemic inflammation (i.e. INFLA-score) as expected (β=-0.10 SD, p<0.001, Rpartial²=0.030), increased high-effort choices (OR=1.32, p=0.044), and marginally decreased effort-related dmPFC signal (β=-0.13, p=0.053, Rpartial²=0.037). These findings show a causal role for inflammation in choosing convenience foods in obesity via increased effort aversion and associated dmPFC processing.

## Introduction

Obesity is a major health problem in the world, affecting one in eight people^1^. Obesity develops from a sustained positive energy balance, in which energy intake exceeds energy expenditure. This imbalance is largely due to maladaptive eating behaviour, such as more frequently choosing for high caloric. easily available foods^2^. Food choices are a form of effort-based decision making, in which one decides whether a certain food is worth exerting a certain amount of effort for, such as cooking, spending money, or chewing^3,4^. The outcomes of these decisions are related to one’s sensitivity to both effort and reward.

Behavioural studies in humans show that obesity is related to altered effort-based decision making. However, findings are inconsistent: some studies report that individuals with obesity were more willing to expend effort for food rewards compared to healthy individuals^5–8^, whereas others were less willing to exert effort^9–11^. Notably, these studies could not disentangle effort from reward, as higher effort was always offered with a higher reward. Therefore, high reward sensitivity in some and effort aversion in others might explain these mixed findings.

Another piece of the puzzle that might explain these paradoxical findings is the presence of chronic low-grade inflammation in obesity. Approximately 50-60% of women and 30-40% of men with obesity show markers of low-grade systemic inflammation^12–15^. This increased inflammatory state originates mainly from (expanding) adipose tissue^16,17^. Additionally, a diet rich in saturated fats, trans fats and foods with a high glycaemic load can contribute to this inflammatory state^18–20^. Adipose tissue expansion triggers the activation and infiltration of macrophages, which promotes the activation of various pro-inflammatory signalling pathways, including the NLRP3 inflammasome and the production of pro-inflammatory cytokines and acute phase proteins^16,21^. Although this inflammatory response is less profound than an acute inflammatory response, it disrupts metabolic pathways in the whole body, including the brain^17^.

In non-obese populations, inflammation has been associated with altered effort-based decision making, particularly with increased aversion to effort. Acute systemic inflammation typically induces sickness behaviour, characterized by low motivation, anhedonia, apathy and fatigue^22–24^. In healthy volunteers, inducing acute inflammation with lipopolysaccharide resulted in increased effort aversion during decision making, without affecting reward sensitivity^25,26^. In addition, systemic inflammation has been associated with altered brain responses in areas that are also affected in obesity and are associated with motivational behaviour, such as mesolimbic areas and the anterior cingulate cortex in the dorsomedial prefrontal cortex (dmPFC)^27,28^. These findings suggest that inflammation can modulate motivation and effort-based decision making, potentially through its effects on effort aversion and brain areas that overlap with those implicated in obesity.

In obesity specifically, the effect of low-grade inflammation on motivation and related brain responses remains unclear. Inflammation has been related to altered responses in the prefrontal cortex, poorer executive functioning, and reduced volume of several cortical and subcortical brain areas in obesity^29–32^. However, studies specifically investigating the effect of inflammation on effort-based decision making, motivation or eating behaviour in obesity are lacking.

Here, we investigated the role of low-grade inflammation on food-related effort-based decision making in brain and behaviour in obesity by (1) studying the association between circulating inflammatory biomarkers and effort and reward sensitivity in individuals with overweight or obesity, and (2) by studying the effect of reducing chronic inflammation using colchicine vs. placebo on effort and reward sensitivity in individuals with obesity. We hypothesized that inflammation would be related to increased aversion to effort and corresponding alterations in effort-related responses in the dmPFC, and that use of colchicine would reverse this effect (preregistered at OSF: https://osf.io/2c3ax).

## Results

### Demographics

We employed a cross-sectional study, including 18-59 year old women with a BMI >27 kg/m^2^, and a randomized controlled trial, including 18-59-year-old women with a BMI >30 kg/m^2^ and low-grade inflammation (CRP >3 mg/L).

In total, 150 participants were included in the cross-sectional fMRI study (BMI >27 kg/m^2^). We excluded 19 participants due to missing immune variables, poor task performance or poor MRI quality, resulting in a total study sample of n=131 (Supplementary Fig. 1).

In the randomized controlled trial, participants received a placebo or the anti-inflammatory drug colchicine, aimed at lowering obesity-related inflammation by inhibition of NLRP3 inflammasome activation and the migration and activation of neutrophils^33–35^. In this study, we included 59 participants (BMI >30 kg/m^2^; CRP >3 mg/L), of which 45 also participated in the cross-sectional study. We excluded 10 participants due to poor task performance or MRI quality, resulting in a total study sample of 49, of which 24 in the colchicine and 25 in the placebo group (Supplementary Fig. 1). Mean compliance was high in both groups (colchicine: 96.8%, placebo: 96.1%). Both treatments were tolerated well, with no serious adverse events; a full list of adverse events is provided in Supplementary Table 1.

Table 1 shows the demographic (baseline) characteristics of both study populations. At baseline, participants in the colchicine group had a higher neutrophil-lymphocyte ratio compared to the placebo group (mean±standard deviation (SD): 2.25±0.65 vs 1.82±0.62, respectively, p=0.021). The groups did not differ on any other demographic, inflammatory or metabolic characteristics.

**Table 1:** Baseline characteristics. ^a^Difference between intervention groups at baseline was tested by independent t-test for normally distributed continuous variables, Wilcoxon-test for skewed continuous variables, and chi-square test for categorical variables. ^b^Median (IQR) is reported due to skewed distribution of the variable. BMI, body mass index; HOMA-IR, Homeostasis Model Assessment of Insulin Resistance; IL, interleukin; ISCED, International Standard Classification of Education; POMS, Profile of Mood States. *P<0.05.

|  | Cross-sectional study (n=131) | Intervention study (n=49) |  | *P-value (colchicine vs placebo)<sup>a</sup>* |
| --- | --- | --- | --- | --- |
|  |  | Colchicine group (n=24) | Placebo group (n=25) |  |
|  | *Mean ± SD or n (%)* | *Mean ± SD or n (%)* | *Mean ± SD or n (%)* |  |
| Age (years)<sup>b</sup> | 47.0 (17.5) | 47.0 (15.8) | 49.0 (15.0) | 0.406 |
| BMI (kg/m<sup>2</sup>) | 35.4 ± 4.2 | 37.2 ± 3.8 | 38.4 ± 4.5 | 0.312 |
| Weight (kg) | 101.5 ± 13.7 | 107.1 ± 11.4 | 112.7 ± 13.2 | 0.117 |
| Waist-to-hip ratio | 0.89 ± 0.09 | 0.89 ± 0.06 | 0.88 ± 0.05 | 0.762 |
| Ethnicity (% Dutch) | 125 (95) | 24 (100) | 25 (100) | - |
| Education (ISCED score) | 3.52 ± 1.39 | 3.33 ± 1.49 | 3.32 ± 1.28 | 0.973 |
| Hormonal birth control (%) | 46 (35) | 10 (42) | 10 (40) | 0.906 |
| Systolic Blood Pressure (mmHg) | 128 ± 15 | 131 ± 16 | 130 ± 11 | 0.710 |
| Diastolic Blood Pressure (mmHg) | 74 ± 10 | 75 ± 10 | 76 ± 10 | 0.691 |
| C-reactive protein (mg/L)<sup>b</sup> | 2.9 (5.1) | 7.6 (9.8) | 5.6 (4.4) | 0.435 |
| White blood cell count (*10<sup>3</sup>/mL) | 6.07 ± 1.35 | 6.62 ± 1.58 | 6.27 ± 1.66 | 0.455 |
| Neutrophil-lymphocyte ratio | 2.01 ± 0.67 | 2.25 ± 0.65 | 1.82 ± 0.62 | 0.021* |
| Neutrophil count (*10<sup>3</sup>/mL) | 3.58 ± 1.03 | 4.03 ± 1.00 | 3.52 ± 1.14 | 0.101 |
| Lymphocytes count (*10<sup>3</sup>/mL) | 1.88 ± 0.51 | 1.94 ± 0.74 | 2.08 ± 0.74 | 0.494 |
| Platelets count (*10<sup>3</sup>/mL) | 272 ± 55 | 282 ± 64 | 280 ± 49 | 0.922 |
| INFLA score (SD) | N.a. | 0.08 ± 0.65 | -0.13 ± 0.50 | 0.210 |
| IL-6 (pg/mL)<sup>b</sup> | 1.26 (1.09) | 2.06 (0.84) | 1.91 (1.02) | 0.404 |
| Adiponectin (µg/mL) | 6.26 ± 3.52 | 5.42 ± 2.79 | 4.17 ± 2.23 | 0.092 |
| Leptin (ng/ml) | 93.7 ± 43.0 | 87.6 ± 37.3 | 102.0 ± 48.7 | 0.252 |
| HbA1c (mmol/mol) | 36.7 ± 3.7 | 37.5 ± 4.0 | 36.7 ± 4.0 | 0.469 |
| Insulin, fasting (mE/L)<sup>b</sup> | 12.6 (9.6) | 15.1 (10.1) | 18.1 (11.8) | 0.290 |
| Glucose, fasting (mmol/L) | 5.35 ± 0.64 | 5.39 ± 0.67 | 5.48 ± 0.60 | 0.631 |
| HOMA-IR | 3.43 ± 2.04 | 3.46 ± 2.02 | 4.67 ± 2.61 | 0.117 |
| Fatigue (POMS-score, range 0-24)<sup>b</sup> | 5.0 (8.0) | 6.5 (8.3) | 5.0 (5.0) | 0.307 |
| Depressive mood (POMS-score, range 0-32)<sup>b</sup> | 2.0 (7.0) | 3.0 (8.0) | 1.0 (7.0) | 0.542 |

### Systemic inflammation

We calculated the INFLA score as composite score of low-grade inflammation, which includes CRP, WBC, platelets and neutrophil/lymphocyte ratio^36^. In the cross-sectional study, we only excluded participants who had an acute infection <4 weeks prior to participation. As expected, higher BMI related to higher levels of inflammation as indicated by the INFLA score (β=0.03 SD, 95% confidence interval (CI): 0.01 to 0.05, p=0.015; Rpartial²=0.045 ;Fig. 1a). In the intervention study, we found that colchicine significantly lowered inflammation, as reflected in the INFLA score, compared with placebo (Time*Group: β=-0.10 SD, 95% CI: -0.15 to -0.04, p<0.001; Rpartial²=0.030; Fig. 1b). Colchicine did not affect BMI compared with placebo (Time*Group: β=0.05, 95% CI: -0.08 to 0.17, p=0.448; Rpartial²=0.000). The effect of the intervention on other immune, metabolic, mood, and anthropometric measures can be found in Supplementary Table 2.

**Fig. 1:**
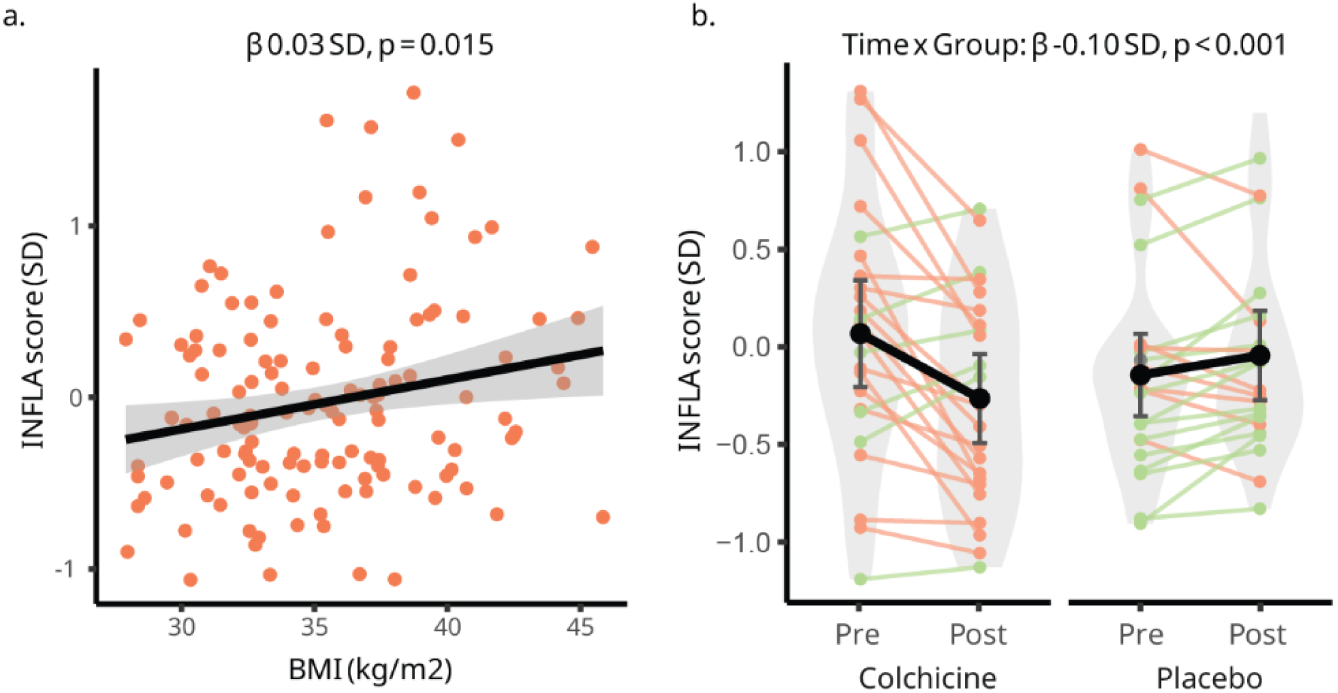
The association between BMI and inflammation, and the effect of colchicine treatment on inflammation. a. The association between BMI and the INFLA score, tested by linear regression modelling. The grey area illustrates the 95% confidence interval. b. Mean(95% confidence interval) change (post – pre) in INFLA-scores split by group. Group difference was tested with mixed linear regression. Green lines indicate an increase over time, whereas orange lines indicate a decrease. BMI, body mass index.

### Main task effects

Participants performed an effort-based decision-making task, designed to disentangle effort and reward sensitivity, during functional MRI^37,38^ (Fig. 2a).

**Fig. 2:**
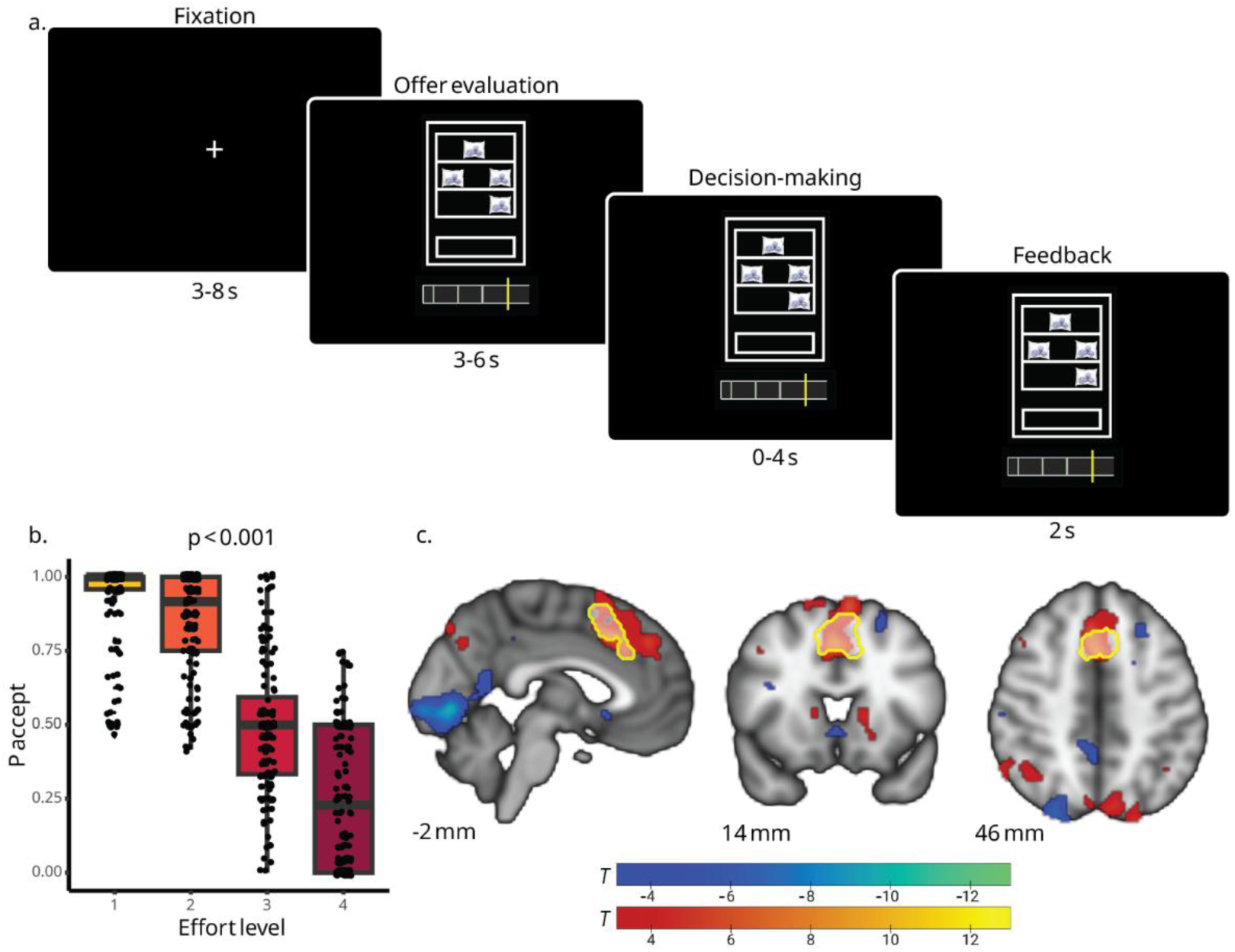
Task design, and behavioural and whole-brain main task effect for effort aversion (n=148). a. During the decision phase of the task, participants saw offers varying in effort level (1-4). On each trial, participants decided whether they were willing to exert a certain amount of effort (squeeze handgrip device; 4 levels) for food rewards that varied in quantity (1 or 4 food items) and calories (high or low caloric). b. Median trial acceptance rates according to effort level, tested using mixed binomial regression analysis. c. Whole-brain task effects for effort are shown in MNI coordinate space at p<0.001 (uncorrected) for illustration purposes. Significant FWE-corrected clusters (p<0.05) are reported in Supplementary Table 4. The region of interest for effort in the medial frontal cortex is outlined in yellow.

To check whether the task manipulation worked as anticipated, we assessed the main task effects across baseline data of all participants in both studies. Mixed binomial regression analysis revealed a main task effect for effort (Fig. 2b; Odds ratio (OR)=0.007, 95% CI: 0.004 to 0.012, p<0.001), showing that participants indeed accepted fewer offers with increasing effort levels. The full regression model can be found in Supplementary Table 3.

Fig. 2c shows the results of the main task effects on BOLD signal across baseline data of all participants in both studies. With increasing effort levels, we found greater BOLD responses in a dmPFC region overlapping with our region-of-interest (ROI) for effort signal, which was defined a-priori based on a meta-analysis of effort-based decision-making tasks^39^ (Fig. 2a; p<0.05 family-wise error (FWE)-corrected). All significant (p<0.05 FWE-corrected) whole brain effort-related BOLD-clusters can be found in Supplementary Table 4.

The intervention groups did not differ at baseline in terms of overall task performance and product wanting measures (Supplementary Table 5). Main task effects for reward quantity and calories are reported in Supplementary Table 3 and 4.

### Inflammation and effort aversion

To assess the effects of inflammation on effort aversion, we investigated how the INFLA score and colchicine treatment were related to acceptance rates of offers with increasing effort levels, while correcting for age and BMI. A higher INFLA score was related to more effort aversion (Fig. 3a; INFLA*Effort: OR=0.27, 95% CI: 0.12 to 0.67, p=0.004), i.e. lower acceptance rates for high effort offers, in the cross-sectional study sample. We performed an ROI analysis to study the association between the INFLA score and effort-related BOLD signal in the dmPFC. This ROI analysis revealed that a higher INFLA score was also associated with stronger effort-related signal in the dmPFC (Fig. 3b; β=0.23 a.u., 95% CI: 0.03 to 0.44, p=0.025, Rpartial²=0.039). An exploratory whole-brain analysis showed that the INFLA score was related to increased effort-related BOLD response in the left posterior cingulum (peak MNI coordinates: 2, -36, 28; k=91; T=4.55; FWE-corrected p(cluster)=0.037). Furthermore, we explored the relationship between BMI and effort aversion, and whether this relationship was explained by inflammation by conducting a mediation analysis. Without controlling for inflammation, BMI was not related to effort aversion (BMI*Effort OR=0.70, 95% CI: 0.44 to 1.13, p=0.144), but was marginally related to increased effort-related BOLD response in de dmPFC (β=0.10, 95% CI: -0.01 to -0.22, p=0.076, Rpartial²=0.024). We found that this relationship between BMI and effort-related signal in the dmPFC was fully mediated by the INFLA score (Fig. 3c; indirect effect: a*b=0.01, p=0.047).

**Fig. 3:**
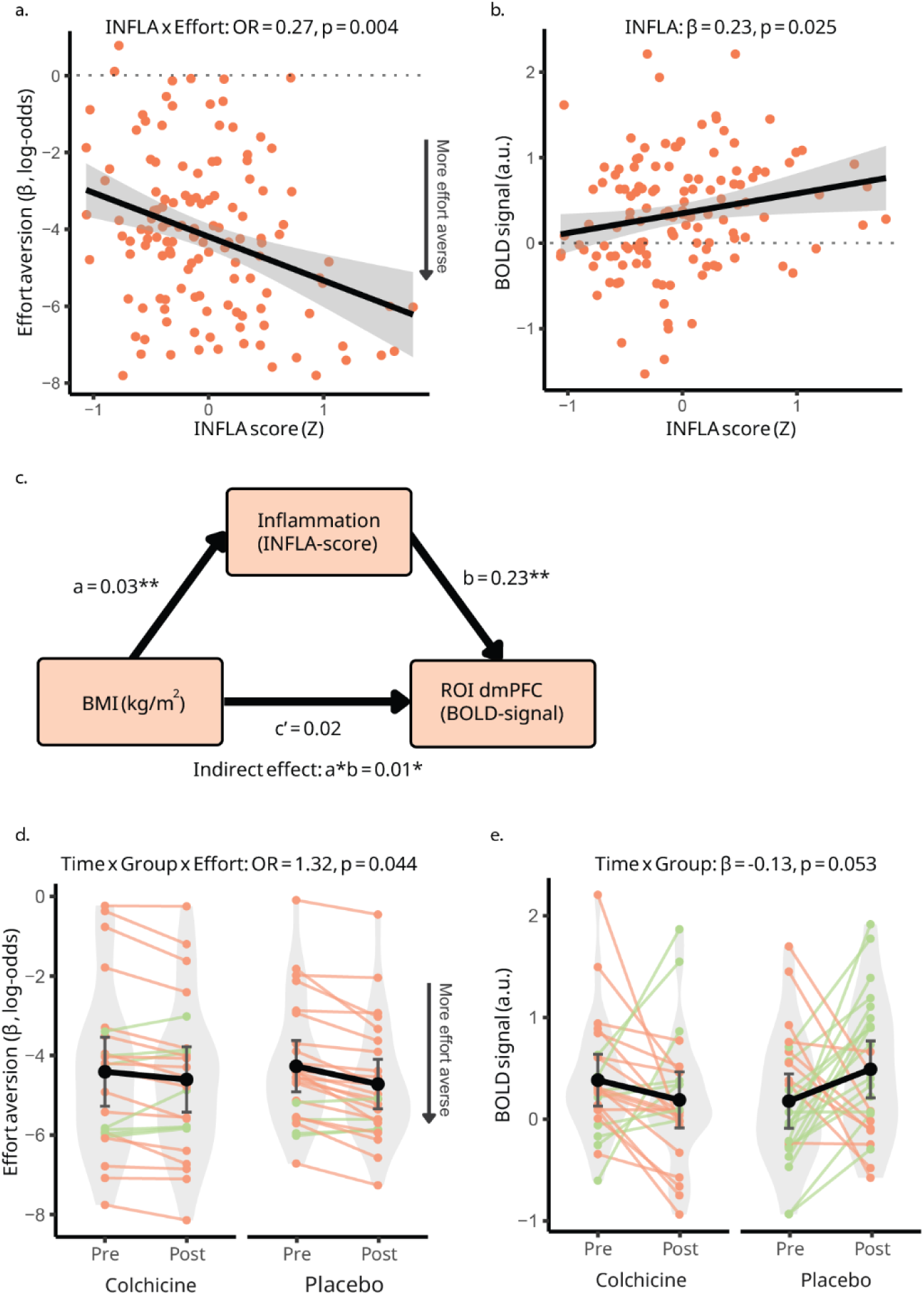
The association between the INFLA score and effort aversion, as well as the effect of colchicine treatment on effort aversion. a. The relationship between the INFLA score and effort aversion, tested by binomial mixed regression modelling. A more negative b for effort aversion on the y-axis illustrates more effort aversion. b. The association between the INFLA score and effort-related blood-oxygen-level-dependent (BOLD) signal, tested by linear regression modelling. c. Mediation analysis to assess the mediating effect of inflammation on the relationship between BMI and effort-related dmPFC BOLD-signal. *P<0.05, **P<0.01. d. Average±standard error change (post – pre) in behavioural betas for effort aversion, tested by mixed linear regression. A more negative b for effort aversion on the y-axis illustrates an increase in effort aversion. d. Average±standard error change (post – pre) in effort-related dmPFC BOLD signal according to group, tested by mixed linear regression. Green lines indicate an increase over time, whereas orange lines indicate a decrease. BMI, body mass index; BOLD, blood-oxygen-level-dependent; dmPFC, dorsomedial prefrontal cortex; OR, odds-ratio.

In the intervention study, we found that colchicine significantly lowered effort aversion compared with placebo, i.e. higher acceptance rates for high-effort offers in the colchicine group (Fig. 3d; Group*Timepoint *Effort: OR=1.32, 95% CI: 1.01 to 1.75, p=0.044). Furthermore, we found a marginal decrease in effort-related signal in the dmPFC in the colchicine group compared with the placebo group (Fig. 3e; β=-0.13 a.u., 95% CI: -0.26 to 0.00, p=0.053, Rpartial²=0.037). From Fig. 3d and 3e, it seems that this effect is driven by colchicine protecting against a further increase in effort aversion and effort-related BOLD-signal the second time the participants performed the task (i.e. post-intervention). The full regression models can be found in Supplementary Table 6 and 7.

### Inflammation and reward quantity and calorie sensitivity

With respect to calorie and reward quantity sensitivity, we found that colchicine increased behavioural sensitivity to calories (Group*Timepoint *Calories: OR=1.16, 95% CI: 1.00 to 1.34, p=0.047). This effect was not paralleled by a correlation with INFLA score in behaviour, by any effects on reward quantity, or by calorie- and reward quantity-related whole-brain effects (see Supplementary Results + Supplementary Discussion).

## Discussion

We investigated the role of inflammation in food-related effort-based decision making in overweight and obesity. In the cross-sectional study, we found that increased inflammation was related to more effort-averse behaviour, and to more effort-related fMRI signal in the dmPFC during a food-related decision-making task. A similar marginal relationship between BMI and effort-related fMRI signal in dmPFC could be completely explained by inflammation. Accordingly, in a randomized controlled trial, we found that colchicine treatment decreased effort-averse behaviour and also marginally decreased effort-related fMRI signal in the dmPFC during the decision-making task.

The fact that all behavioural effects pointed in the same direction in this multi-level approach, combining behavioural and neuroimaging measures in both a cross-sectional and a pharmacological intervention design, strengthens the confidence in the robustness of these correlations and causal effects of inflammation on effort aversion. We did not observe similarly consistent effects for reward sensitivity in our exploratory analyses (calories or quantity; see Supplementary Discussion).

Evidence regarding the role of inflammation in eating behaviour in human obesity is still scarce. Preclinical evidence, however, supports the role of inflammation in modulating eating behaviour. In rodents, pharmacological inhibition of the NLRP3 inflammasome—the same target we blocked using colchicine—decreased both food intake and body fat mass in mice with diet-induced obesity, with effect sizes comparable to the weight-loss medication semaglutide^40^. Furthermore, also other types of anti-inflammatory treatment have been shown to reduce weight gain and/or food intake in high-fat diet induced obesity in rodents ^41–43^. In humans, studies have investigated food choice behaviour in obesity but largely without considering the inflammatory status of participants. Findings of these studies are inconsistent, with some reporting reduced effort expenditure^9–11^ and others reporting increased effort expenditure for food rewards^5–8^. Our results, in combination with these preclinical findings, indicate that there may be two subgroups in obesity based on their inflammatory level: in individuals with higher low-grade inflammation, altered food choice behaviour could be primarily driven by effort aversion, as we show currently, whereas in those without low-grade inflammation, other mechanisms, such as increased reward sensitivity due to genetic predisposition^44,45^, may play an important role. Future studies investigating eating behaviour in obesity should therefore aim to take into account the inflammatory status of participants.

Our results confirm our hypothesis that low-grade inflammation leads to effort aversion and increased effort-related signal in the dmPFC in obesity. The results are in line with a recent study that employed a similar anti-inflammatory design, decision-making task, and a-priori ROI in a depressed population with low-grade inflammation^46^. There, the anti-inflammatory drug infliximab also lowered effort aversion during an effort-based decision-making task compared to placebo. Additionally, while they did not find a difference in dmPFC responses between groups, they found that a decrease in the pro-inflammatory marker TNF was related to a decrease in activity in the dmPFC. Furthermore, in healthy participants, LPS-induced acute inflammation decreased acceptance of high-effort offers but did not affect the participants’ sensitivity to rewards^25,26^. Together, these findings support a link between inflammation and effort aversion. Our work extends these findings by demonstrating this effect in obesity, in the context of food rather than monetary rewards, using a unique combined cross-sectional and randomized intervention design.

Inflammation affects effort sensitivity likely via disruption of dopamine signalling. Obesity-related inflammation mainly originates from enlarged fat cells in expanding adipose tissue, which attracts macrophages and leads to secretion of cytokines^16,17^. In addition, certain nutrients in the diet, such as high intake of saturated fats, can directly and indirectly induce an inflammatory response by activating immune pathways such as the NF-kB pathways and by increasing gut permeability^47^. Pro-inflammatory cytokines can travel through the blood-brain barrier due to increased permeability during inflammation^48^. Here, cytokines such as IL-6 and TNF-alpha impair – amongst others - dopamine synthesis, release and availability by directly lowering levels of cofactors such as BH4, which is needed for the formation of the precursor L-DOPA from phenylalanine and tyrosine^49^. As dopamine modulates activity in brain areas related to motivation and decision making, such as the prefrontal cortex and striatum, impaired dopamine neurotransmission can lead to altered motivational behaviour, decision making, and control of food intake^50^. Future studies should clarify the role of dopamine in inflammation-related changes in effort aversion and motivational behaviour.

The inclusion of only women limits the ability to draw conclusions about the inflammatory effects on decision making in men. We only included women because of their higher susceptibility to overweight and obesity, obesity-related inflammation, and depression, which is characterized by motivational deficits^12,51,52^. For instance, in men, an inflammatory challenge did not affect reward sensitivity, while it did in women^53^. Differences in inflammation-related pathways, fat distribution, hormonal regulation and behavioural responses may lead to these distinct associations between inflammation and decision making^54–56^. Future research should therefore include both sexes to better understand potential sex-specific mechanisms.

In addition, while our behavioural task allowed us to elucidate the mechanism behind food-related decision making under controlled conditions, it does not fully reflect the complexity of real-life food choice. Real-life food choice involves other environmental factors, such as social context, hunger, and availability which are not easily captured in a laboratory paradigm. Future studies should investigate how these mechanistic findings relate to more ecologically valid measures, such as real-life food choice, motivational behaviour and adherence to lifestyle interventions.

In conclusion, our findings show that low-grade inflammation in obesity increases aversion to effort during food-related decision making, and increases effort-related brain-activity in the dmPFC. Importantly, anti-inflammatory treatment reversed this effect both in the behaviour and, marginally, in the neural signal. This demonstrates a causal role of inflammation in promoting convenient, fast-food choices in obesity. These findings highlight the role of inflammation as a potential target for the treatment of obesity, for example through the implementation of anti-inflammatory strategies into existing lifestyle programs.

## Methods

### Population and design

This functional MRI study had a combined cross-sectional and intervention design (Fig. 4). Participants for both parts of the study were recruited from advertisement on social media and in local newspapers, and the Radboud University’s participant databases. Additionally, participants from the intervention study were recruited after participation in the cross-sectional study. In the cross-sectional study, we included 18-59-year-old women with a BMI >27 kg/m^2^. In the intervention study, we included 18-59-year-old women with a BMI >30 kg/m^2^ and low-grade inflammation as indicated by CRP >3 mg/L. For both studies, we only included women without diabetes, autoimmune, psychiatric or inflammatory disease, who had not been sick or vaccinated at least 4 weeks before participation, and did not have CRP levels indicating acute inflammation (i.e., excluding CRP>10 mg/L for BMI ≤31 kg/m^2^ and CRP>22.1 mg/L for BMI >31 kg/m^2^)^57^. The full list of in- and exclusion criteria of both studies can found in Supplementary Methods 1.

**Fig. 4:**
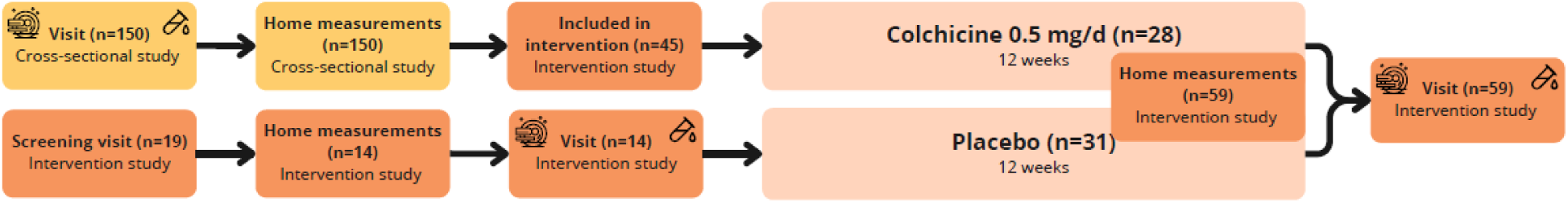
Study design. The cross-sectional study is illustrated in yellow, and the intervention study in orange.

For the cross-sectional study, participants fasted overnight before coming to the test centre, where we took a blood sample, gave them a standardized breakfast, and performed behavioural and neuroimaging measurements. The full description of the procedures can be found in Supplementary Methods 2).

The intervention study had a randomized double-blind, placebo-controlled trial design (ClinicalTrials.gov identifier: NCT05785429). Participants were randomized 1:1 to receive one capsule of 0.5 mg colchicine or placebo daily for 12 weeks. Colchicine lowers inflammation by blocking the NLRP3 inflammasome, and has been shown to lower inflammation in obesity before^33,34^. Randomization was done using computer-generated randomization sequence implemented in Castor Electronic Data Capture (Castor EDC) and was stratified by baseline age (strata: 18-40, 41-59 years), BMI (strata: 30-35, >35 kg/m^2^) and CRP (strata: 3.0-6.5 mg/L, >6.5 mg/L) to ensure balanced distribution of these variables across groups. Colchicine (Centrafarm B.V, Etten-Leur, the Netherlands) and placebo capsules were stored and dispensed by the clinical pharmacy of the Radboudumc, Nijmegen, the Netherlands, and were identical in appearance, taste and packaging. Baseline and follow-up blood, neuroimaging, and behavioural assessments were conducted in identical test days as for the cross-sectional study. For participants enrolled in both studies, their cross-sectional assessments served as baseline measurements. Compliance was monitored by capsule count at the follow-up visit.

Both studies were approved by the Medical Ethics Review Committee (METC) Oost-Nederland, Nijmegen, the Netherlands (reference number of cross-sectional study: NL77503.091.21 and intervention study: NL79408.091.21). All procedures were conducted in accordance with the Declaration of Helsinki and written informed consent was obtained from all participants prior to participation.

### Inflammatory markers

We measured wide-range CRP (0.5-200 mg/L) via a capillary blood sample collected by finger prick (QuikRead go®, Aidian). Furthermore, venous blood samples were collected and processed within 1h of collection. Blood cell counts (white blood cell count (WBC), lymphocytes, neutrophils and platelets) were measured in whole blood using Sysmex XN-450 Hematology Analyzer (Sysmex Corp., Kobe, Japan). Adiponectin and leptin were measured in EDTA-plasma using solid-phase sandwich ELISA (Quantikine ELISA Kit, R&D Systems, Minneapolis, USA). Glucose was measured in sodium fluoride/Na2EDTA-plasma using a Biosen C-Line; EKF Diagnostics glucose analyzer. Plasma IL-6 concentrations were measured with high sensitivity by electrochemiluminescence using a MesoScale Discovery (MSD) multiplex system and a validated Vplex-human proinflammatory panel I (MesoScale Discovery, Rockville, Maryland, USA). HbA1c concentrations were determined by ion-exchange high-performance liquid chromatography (HPLC) on the Tosoh G11 analyzer (Tosoh Bioscience, Tokyo, Japan). Serum insulin concentrations were measured on the Roche Cobas analyser (module e801) using an electrochemiluminescence immunoassay (ECLIA) (Roche Diagnostics, Mannheim, Germany).

### Experimental task: effort-based decision making

We measured effort aversion with an adapted version of an existing effort-based decision-making task^37,38^. Participants decided whether a food reward (varying in quantity and calories) was worth a certain level of effort (squeezing in a handgrip device). The task was divided in three phases: (1) calibration of and familiarization with the effort levels outside the scanner, (2) decision-making in the MRI scanner, and (3) execution outside the scanner. By separating the decisions from the execution of the trials, we minimized fatigue effects on the participant’s decisions.

In the calibration phase, the participant’s maximum voluntary contraction (MVC) was measured three times for 5s. The best attempt was used to define four effort levels (10%, 33%, 56%, 80% MVC). They practiced each level three times to get familiar with the level’s required effort. Participants then chose whether they preferred sweet (chocolates and blueberries) or savoury foods (crisps and cucumber), and we asked them to rate their wanting of these food items on a 100mm VAS scale. Their preferred type of food was then used throughout the task as reward.

The decision phase was performed inside the MRI scanner (Fig. 2a). On each trial, participants decided whether to exert one of the four effort levels to obtain food rewards varying in quantity (1 or 4 items) and caloric content (low vs. high). Sweet-preferring participants received blueberries as low-caloric and chocolates as high-caloric option, while savoury-preferring participants received cucumber as low-caloric and crisps as high-caloric option. Offers were visualized as a vending machine holding the offered food items, and a bar indicating the required effort-level. Each offer was presented for 3–6s, after which

Yes/No options appeared randomly on the left and right side of the screen. Participants responded within 4s using a button box, after which their choice was shown (2s) followed by a fixation cross (3–8s). All combinations of effort × reward quantity × calorie level were presented six times (96 trials total), randomized into six blocks of 16 trials with 20s fixation breaks in between. The total duration of the decision phase was 22m.

In the execution phase outside the scanner, participants were presented with 32 of their previous choices. Accepted offers required squeezing the handgrip to earn the food, while rejected offers were followed by a 5s wait. At the end of the session, participants received the snacks they earned.

### MRI acquisition

The MRI scans were performed on a 3.0 Tesla Siemens MAGNETOM Skyra MRI scanner, using a 32-channel head coil. During each visit, a structural whole-brain scan was acquired using a T1-weighted MP2RAGE sequence (repetition time: 6000 ms, echo time: 2.34 ms, 6* flip angle, FOV: 256x256 matrix, 176 sagittal slices, voxel size: 1.0x1.0x1.0 mm, 1.0 mm slice thickness). The functional scans measured blood oxygen level dependent (BOLD) signals by using a T2*-weighted gradient echo single-echo multi-band 4 echo planar imaging (EPI) sequence (repetition time: 1500 ms, echo time: 33.40 ms, 75**°** flip angle, FOV: 213x213 matrix, 68 slices, interleaved acquisition, voxel size: 2.0x2.0x2.0 mm, 2.0 mm slice thickness).

The functional MRI data were pre-processed using *fMRIPrep* 24.0.0 (Esteban et al. (2019); Esteban et al. (2018); RRID:SCR_016216), which is based on *Nipype* 1.8.6 (K. Gorgolewski et al. (2011); K. J. Gorgolewski et al. (2018); RRID:SCR_002502). The full pre-processing pipeline is described in Supplementary Methods 3. We used SPM12 (Welcome Trust Centre for Neuroimaging, University College London, UK) running in Matlab R2024b (Mathworks, Inc., MA, United States) to spatially smooth the final pre-processed data with a 6mm FWHM kernel.

## Statistics

### Data quality checks

We excluded participants who accepted more than 95% or less than 5% of the offers, and who responded with the same button in more than 70% of the trials, as they may not have properly understood or engaged with the task. Furthermore, we excluded participants with bad MRI data quality due to acquisition problems or with >0.5 mm average framewise displacement during the task.

Continuous variables were screened for outliers using >3 SD cut-off and visual inspection of histograms and boxplots. Data points were excluded if the values reflected a measurement error or were biologically implausible. MRI analyses were conducted in SPM12 (Welcome Trust Centre for Neuroimaging, University College London, UK) running in Matlab R2024b (Mathworks, Inc., MA, United States). All other statistical analyses were performed in RStudio (version 2025.05.1).

### Demographics

We calculated mean±SD for continuous and n (%) for categorical baseline variables of both the cross-sectional study population and the two intervention groups. We tested the baseline differences between the two intervention groups using independent t-tests and chi-square tests. In the cross-sectional study, we investigated the relationship between BMI and the INFLA score using linear regression modelling.

In the intervention study, we tested the effect of the intervention on immune, metabolic and anthropometric variables using mixed linear regression with the immune, metabolic or anthropometric variable as outcome, Time (baseline, follow-up) as random factor, and Time, Group and its interaction as fixed factors of interest.

### Effect of intervention on inflammation

As inflammation measure, we used the INFLA score, a composite score of CRP, WBC, platelets and neutrophil/lymphocyte ratio commonly used in low-grade inflammation research^36^. The INFLA score was computed by averaging the Z-scores of these four variables. We tested the effect of the intervention on the INFLA score using mixed linear regression with the INFLA score as outcome, Time as random factor, and Time, Group and its interaction as fixed factors of interest.

### Behavioural analysis: main task effects

To assess the main task effects, we combined baseline data from both the cross-sectional study and the intervention study. To study the effects of effort, reward quantity, and calories on offer acceptance (yes, no), we executed a mixed binomial regression analysis, in which we included Effort, RewardQuantity, and Calories and its interactions as both fixed and random factors.

### Behavioural analysis: relationship with inflammation

In the cross-sectional study, we tested whether the INFLA score was related to effort aversion, reward quantity and calorie sensitivity by using binomial mixed regression modelling with offer acceptance (yes/no) as outcome variable, Effort, RewardQuantity, Calories, and its interactions as random factors, and INFLA, Effort, Calories, RewardQuantity, and its interactions as fixed factors. We adjusted the model by adding Age and BMI as fixed factors of non-interest. We Z-scored all continuous predictors in the model.

Furthermore, we explored the relationship between BMI and effort aversion, and if significant, a potential mediating effect of inflammation on this relationship using structural equation modelling (SEM) using the Lavaan package (version 0.6.19)^58^.

### Behavioural analysis: effect of intervention

We tested the effect of the intervention on effort aversion, reward quantity and calorie sensitivity by using binomial mixed regression modelling with offer acceptance (yes/no) as outcome variable, Effort, RewardQuantity, and Calories, Timepoint, and its interactions as random factors, Effort, RewardQuantity, and Calories, Timepoint, Group, and its interactions as fixed factors of interest. We adjusted the model by adding baseline Age, BMI and CRP as fixed factors of non-interest. We Z-scored all continuous predictors in the model.

### Functional MRI analysis: main task effects

To study the main task effects on BOLD signal, we combined fMRI baseline data from the cross-sectional and intervention study. For each participant, we constructed a first-level generalized linear model (GLM) with four regressors reflecting the 2x2 design of the task for Calories (high vs. low) and Reward Quantity (high vs. low). Effort was modelled as parametric modulator for each regressor, resulting in four additional regressors. All regressors were convolved with the SPM-standard canonical hemodynamic response function (HRF). We included six realignment parameters, framewise displacement, the first 5 aCompCor and tCompCor noise regressors, and ICA-AROMA noise regressors (derived from fmriprep, see Supplementary Methods 3).

To assess effect of effort, calories and reward quantity on event-related BOLD signal, we computed three contrast images for each participant using paired t-tests. For the effort contrast we summed the four parametric modulators. For the calories contrast, we subtracted the low-calorie regressors from the high-calorie regressors. For the reward quantity regressors, we subtracted the low-reward regressors from the high-reward regressors.

At the group level, we studied the main effects of effort, reward quantity- and calories on whole-brain BOLD signal using one-sample t-tests on the corresponding first-level contrast images. We used an initial cluster-forming threshold of p<0.001 and considered clusters significant at whole brain level if their FWE-corrected p-value was below 0.05. Cluster sizes and peak coordinates are reported in MNI space.

### Functional MRI analysis: associations with INFLA

To study the association between inflammation and effort-related BOLD signal in the cross-sectional study, we performed a confirmatory region-of-interest (ROI) analysis and additionally explored whole-brain effects.

First, we studied the association between the INFLA score and effort-related BOLD signal in the dmPFC using a pre-defined ROI from a meta-analysis studying effort valuation in the brain^39^ (yellow outline in Fig. 2c). Mean beta values from the effort-contrast were extracted for each participant and analysed using linear regression, including the INFLA score as predictor and Age and BMI as covariates.

Furthermore, we explored the relationship between BMI and effort-related dmPFC responses, and a potential mediating effect of inflammation on this relationship using structural equation modelling (SEM) using the Lavaan package (version 0.6.19)^58^. We defined BMI as the predictor variable, the INFLA score as the mediator, and dmPFC effort-related signal as the outcome variable. We used bootstrapping with 5,000 resamples to estimate indirect effects. We included Age in the model as covariate.

To assess whole-brain associations, we repeated the group-level analyses for Effort, Reward Quantity, and Calories with the INFLA score as covariate of interest, and Age and BMI as covariates of non-interest.

### Functional MRI analysis: effect of the intervention

To study the effect of the intervention on effort-related BOLD signal, we executed a confirmatory ROI analysis and additionally explored whole-brain effects.

First, we studied the difference between groups in effort-related BOLD signal in the dmPFC using the same pre-defined ROI^39^. For each participant, we built first-level GLMs at baseline and follow-up, each with the eight task parameters as described in the main task effect analysis. For each timepoint, effort contrasts were computed by summing the four parametric modulators, and mean ROI betas were extracted. We tested the effect of the intervention on these ROI betas using mixed regression modelling, with ROI betas as outcome, Timepoint as random factor, Time, Group, and Time x Group as fixed factors of interest, and baseline Age, BMI and CRP as fixed factors of non-interest.

To assess whole-brain effects of the intervention, we constructed a first-level GLM including the same eight task parameters for each timepoint. To study the effect on effort-related signal, we subtracted the baseline effort parametric regressors from the follow-up effort parametric regressors. To study the effect on calorie-related signal and reward quantity-related signal, we subtracted the high vs. low calorie/reward quantity related signal at baseline from the calorie/reward quantity related signal at follow-up. At group level, we then performed a voxel-wise GLM analysis using two-sample t-tests to compare the colchicine and placebo groups for the contrasts for effort, reward quantity and calories.

## Acknowledgements

We would like to thank all participants who contributed their time and effort to the FLAIR study. Furthermore, we would like to thank the master’s and bachelor’s students who assisted with the data collection for their valuable practical contributions; Lisa-Katrin Kaufmann for her advice on the methodology and statistical analysis of the study; Manon Dumont, Julia Brake, and Ajie Mandala for performing the immunological analyses and processing the blood samples; Ilke de Lange for her assistance with the data collection; Paul Gaalman for his assistance with the MRI data collection; and Britt Lambregts for het contributions to the code of the behavioural task. This project has received funding from the European Research Council (ERC) under the European Union’s Horizon 2020 research and innovation programme (grant agreement No 852189 to EA).

## Contributions

E.A. conceptualized the project. E.A., R.S., R.B., and J.S. designed the methodology. J.S., L.O. and M.H. collected data. J.S. and R.B. analysed the data. C.M.M. provided medical supervision. J.S. wrote the initial draft, supervised by E.A. and R.B.. All authors reviewed and edited the manuscript.

## Data availability

Data supporting the findings of this study are available from the Radboud Data Repository via the following URL: https://doi.org/10.34973/ahd1-bj58.

## Code availability

Code for all analysis in the manuscript and supplementary material are available from the Radboud Data Repository via the following URL: https://doi.org/10.34973/ahd1-bj58.

## References

1. Boutari, C. & Mantzoros, C. S. A 2022 update on the epidemiology of obesity and a call to action: as its twin COVID-19 pandemic appears to be receding, the obesity and dysmetabolism pandemic continues to rage on. Metabolism 133, 155217 (2022).

2. Romieu, I. et al. Energy balance and obesity: what are the main drivers? Cancer Causes Control 28, 247–258 (2017).

3. Chong, T. T.-J., Bonnelle, V. & Husain, M. Quantifying motivation with effort-based decision-making paradigms in health and disease. in Progress in Brain Research vol. 229 71–100 (Elsevier, 2016).

4. Brassard, S. L. & Balodis, I. M. A review of effort-based decision-making in eating and weight disorders. Progress in Neuro-Psychopharmacology and Biological Psychiatry 110, 110333 (2021).

5. Epstein, L. H. et al. Food Reinforcement, the Dopamine D2 Receptor Genotype, and Energy Intake in Obese and Nonobese Humans. Behav Neurosci 121, 877–886 (2007).

6. Giesen, J. C. A. H., Havermans, R. C., Douven, A., Tekelenburg, M. & Jansen, A. Will Work for Snack Food: The Association of BMI and Snack Reinforcement. Obesity 18, 966–970 (2010).

7. Rollins, B. Y., Loken, E., Savage, J. S. & Birch, L. L. Measurement of food reinforcement in preschool children. Associations with food intake, BMI, and reward sensitivity. Appetite 72, 21–27 (2014).

8. Saelens, B. E. & Epstein, L. H. Reinforcing value of food in obese and non-obese women. Appetite 27, 41–50 (1996).

9. Mata, F. et al. Reduced Willingness to Expend Effort for Reward in Obesity: Link to Adherence to a 3-Month Weight Loss Intervention. Obesity 25, 1676–1681 (2017).

10. Mathar, D., Horstmann, A., Pleger, B., Villringer, A. & Neumann, J. Is it Worth the Effort? Novel Insights into Obesity-Associated Alterations in Cost-Benefit Decision-Making. Frontiers in Behavioral Neuroscience 9, (2016).

11. Miras, A. D. et al. Gastric bypass surgery for obesity decreases the reward value of a sweet-fat stimulus as assessed in a progressive ratio task. Am J Clin Nutr 96, 467–473 (2012).

12. Aronson, D. et al. Obesity is the major determinant of elevated C-reactive protein in subjects with the metabolic syndrome. Int J Obes Relat Metab Disord 28, 674–679 (2004).

13. Festa, A. et al. The relation of body fat mass and distribution to markers of chronic inflammation. Int J Obes Relat Metab Disord 25, 1407–1415 (2001).

14. Marques-Vidal, P. et al. Association between inflammatory and obesity markers in a Swiss population-based sample (CoLaus Study). Obes Facts 5, 734–744 (2012).

15. Visser, M., Bouter, L. M., McQuillan, G. M., Wener, M. H. & Harris, T. B. Elevated C-reactive protein levels in overweight and obese adults. JAMA 282, 2131–2135 (1999).

16. Rohm, T. V., Meier, D. T., Olefsky, J. M. & Donath, M. Y. Inflammation in obesity, diabetes, and related disorders. Immunity 55, 31–55 (2022).

17. Saltiel, A. R. & Olefsky, J. M. Inflammatory mechanisms linking obesity and metabolic disease. J Clin Invest 127, 1–4 (2017).

18. Buyken, A. E. et al. Association between carbohydrate quality and inflammatory markers: systematic review of observational and interventional studies. Am J Clin Nutr 99, 813–833 (2014).

19. Shivappa, N., Steck, S. E., Hurley, T. G., Hussey, J. R. & Hébert, J. R. Designing and developing a literature-derived, population-based dietary inflammatory index. Public Health Nutr 17, 1689–1696 (2014).

20. Zhou, H., Urso, C. J. & Jadeja, V. Saturated Fatty Acids in Obesity-Associated Inflammation. JIR 13, 1–14 (2020).

21. Rheinheimer, J., de Souza, B. M., Cardoso, N. S., Bauer, A. C. & Crispim, D. Current role of the NLRP3 inflammasome on obesity and insulin resistance: A systematic review. Metabolism 74, 1–9 (2017).

22. Dantzer, R. Cytokine-induced sickness behavior: mechanisms and implications. Ann N Y Acad Sci 933, 222–234 (2001).

23. De Marco, R. et al. Inflammation-induced reorientation of reward versus punishment sensitivity is attenuated by minocycline. Brain Behav Immun 111, 320–327 (2023).

24. Lasselin, J. et al. Lipopolysaccharide Alters Motivated Behavior in a Monetary Reward Task: a Randomized Trial. Neuropsychopharmacology 42, 801–810 (2017).

25. Draper, A. et al. Effort but not Reward Sensitivity is Altered by Acute Sickness Induced by Experimental Endotoxemia in Humans. Neuropsychopharmacol. 43, 1107–1118 (2018).

26. Lambregts, B. I. H. M. et al. Fatigue during acute systemic inflammation is associated with reduced mental effort expenditure while task accuracy is preserved. Brain, Behavior, and Immunity 112, 235–245 (2023).

27. García-García, I. et al. Reward processing in obesity, substance addiction and non-substance addiction. Obesity Reviews 15, 853–869 (2014).

28. Kraynak, T. E., Marsland, A. L., Wager, T. D. & Gianaros, P. J. Functional neuroanatomy of peripheral inflammatory physiology: A meta-analysis of human neuroimaging studies. Neuroscience & Biobehavioral Reviews 94, 76–92 (2018).

29. Bao, Y. et al. Chronic Low-Grade Inflammation and Brain Structure in the Middle-Aged and Elderly Adults. Nutrients 16, 2313 (2024).

30. Kaufmann, L.-K. et al. Surgery-induced reduction in inflammation relates to improved neural inhibitory control in obesity. Brain Behav Immun 129, 829–838 (2025).

31. Naomi, R., et al. The Role of Oxidative Stress and Inflammation in Obesity and Its Impact on Cognitive Impairments—A Narrative Review. Antioxidants 12, (2023).

32. Yang, Y. et al. The association between obesity and lower working memory is mediated by inflammation: Findings from a nationally representative dataset of U.S. adults. *Brain*, Behavior, and Immunity 84, 173–179 (2020).

33. Demidowich, A. P., Davis, A. I., Dedhia, N. & Yanovski, J. A. Colchicine to decrease NLRP3-activated inflammation and improve obesity-related metabolic dysregulation. Medical Hypotheses 92, 67–73 (2016).

34. Demidowich, A. P. et al. Colchicine’s effects on metabolic and inflammatory molecules in adults with obesity and metabolic syndrome: results from a pilot randomized controlled trial. Int J Obes 44, 1793–1799 (2020).

35. de Bulhões, F. V. et al. The Action of Colchicine in Patients with Metabolic Syndrome and Obesity: Perspectives and Challenges. Metabolites 14, 629 (2024).

36. Bonaccio, M. et al. A score of low-grade inflammation and risk of mortality: prospective findings from the Moli-sani study. Haematologica 101, 1434–1441 (2016).

37. Bonnelle, V., Manohar, S., Behrens, T. & Husain, M. Individual Differences in Premotor Brain Systems Underlie Behavioral Apathy. Cerebral Cortex 26, 807–819 (2016).

38. Contreras-Huerta, L. S., Lockwood, P. L., Bird, G., Apps, M. A. J. & Crockett, M. J. Prosocial behavior is associated with transdiagnostic markers of affective sensitivity in multiple domains. Emotion 22, 820–835 (2022).

39. Castanheira, K. da S., Spreng, R. N., Vassena, E. & Otto, A. R. The neural basis of cost-benefit trade-offs in effort investment: a quantitative activation likelihood estimation meta-analysis. 2022.10.28.513278 Preprint at 10.1101/2022.10.28.513278 (2022).

40. Thornton, P. et al. Reversal of High Fat Diet-Induced Obesity, Systemic Inflammation, and Astrogliosis by the NLRP3 Inflammasome Inhibitors NT-0249 and NT-0796. J Pharmacol Exp Ther **388**, 813–826 (2024).

41. Alsaggar, M. et al. Silibinin attenuates adipose tissue inflammation and reverses obesity and its complications in diet-induced obesity model in mice. BMC Pharmacol Toxicol 21, 8 (2020).

42. Yang, J., Li, W. & Wang, Y. Capsaicin Reduces Obesity by Reducing Chronic Low-Grade Inflammation. International Journal of Molecular Sciences 25, 8979 (2024).

43. Wu, Y., Yu, Y., Szabo, A., Han, M. & Huang, X.-F. Central Inflammation and Leptin Resistance Are Attenuated by Ginsenoside Rb1 Treatment in Obese Mice Fed a High-Fat Diet. PLoS ONE 9, (2014).

44. Rapuano, K. M. et al. Genetic risk for obesity predicts nucleus accumbens size and responsivity to real-world food cues. Proc Natl Acad Sci U S A 114, 160–165 (2017).

45. Rohde, K. et al. Genetics and epigenetics in obesity. Metabolism 92, 37–50 (2019).

46. Treadway, M. T. et al. A randomized proof-of-mechanism trial of TNF antagonism for motivational deficits and related corticostriatal circuitry in depressed patients with high inflammation. Mol Psychiatry 30, 1407–1417 (2025).

47. Christ, A., Lauterbach, M. & Latz, E. Western Diet and the Immune System: An Inflammatory Connection. Immunity 51, 794–811 (2019).

48. Galea, I. The blood-brain barrier in systemic infection and inflammation. Cell Mol Immunol 18, 2489–2501 (2021).

49. Felger, J. C. The Role of Dopamine in Inflammation-Associated Depression: Mechanisms and Therapeutic Implications. Curr Top Behav Neurosci 31, 199–219 (2017).

50. Wallace, C. W. & Fordahl, S. C. Obesity and dietary fat influence dopamine neurotransmission: exploring the convergence of metabolic state, physiological stress, and inflammation on dopaminergic control of food intake. Nutr Res Rev 35, 236–251 (2022).

51. Eid, R. S., Gobinath, A. R. & Galea, L. A. M. Sex differences in depression: Insights from clinical and preclinical studies. Progress in Neurobiology 176, 86–102 (2019).

52. Kanter, R. & Caballero, B. Global gender disparities in obesity: a review. Adv Nutr 3, 491–498 (2012).

53. Moieni, M. et al. Sex Differences in the Relationship Between Inflammation and Reward Sensitivity: A Randomized Controlled Trial of Endotoxin. Biol Psychiatry Cogn Neurosci Neuroimaging 4, 619–626 (2019).

54. Lewis, C. A., Grahlow, M., Kühnel, A., Derntl, B. & Kroemer, N. B. Women compared with men work harder for small rewards. Sci Rep 13, 5456 (2023).

55. Mauvais-Jarvis, F. Sex differences in metabolic homeostasis, diabetes, and obesity. Biology of Sex Differences 6, 14 (2015).

56. ter Horst, R., et al. Sex-Specific Regulation of Inflammation and Metabolic Syndrome in Obesity. Arterioscler Thromb Vasc Biol 40, 1787–1800 (2020).

57. Ishii, S. et al. Gender, Obesity and Repeated Elevation of C-Reactive Protein: Data from the CARDIA Cohort. PLoS ONE 7, e36062 (2012).

58. lavaan: An R Package for Structural Equation Modeling | Journal of Statistical Software. https://www.jstatsoft.org/article/view/v048i02.

