## Supplementary material for "Low-grade inflammation in obesity causes low-effort food choice"

Supplementary material for: Scholing et al., *Low-grade inflammation in obesity causes low-effort food choice*

### Supplementary Table 1: Adverse events in the intervention study

| | Total Counts | Colchicine Counts | Placebo Counts |
|---|---|---|---|
| Headache | 31 | 16 | 15 |
| Gastrointestinal complaints | 30 | 13 | 17 |
| Cold or flu-like symptoms | 30 | 10 | 20 |
| Fatigue | 8 | 2 | 6 |
| Joint or muscle pain | 18 | 11 | 7 |
| Heavy menstrual flow | 7 | 5 | 2 |
| Other infections | 7 | 3 | 4 |
| Rash | 1 | 0 | 1 |
| Shortness of breath | 1 | 0 | 1 |
| Nerve pain | 1 | 1 | 0 |
| Benign tumor diagnosis | 1 | 1 | 0 |
| Allergic reaction | 1 | 1 | 0 |
| Endometrial polyp | 1 | 1 | 0 |
| Dizziness | 1 | 1 | 0 |
| Restless legs | 1 | 1 | 0 |
| Easy bruising | 1 | 0 | 1 |
| Hot flashes | 1 | 1 | 0 |
| Unusual taste in the mouth | 1 | 0 | 1 |

#### Supplementary Table 2: Intervention effects on anthropometric, immune, metabolic, and mood outcomes

| | Colchicine (n=24) | | Placebo (n=25) | | | | P-value[a] |
|---|---|---|---|---|---|---|---|
| | Baseline | Follow-up | Baseline | Follow-up | Time x Group Effect[a] | | |
| | Mean ± SD | Mean ± SD | Mean ± SD | Mean ± SD | β (95% CI) | $R_{partial}^2$ | |
| BMI (kg/m$^2$) | 37.2 ± 3.8 | 37.3 ± 4.1 | 38.4 ± 4.5 | 38.3 ± 4.8 | 0.1 (-0.1, 0.2) | 0.000 | 0.448 |
| Weight (kg) | 107.1 ± 11.4 | 107.4 ± 12.0 | 112.7 ± 13.2 | 112.4 ± 13.3 | 0.2 (-0.2, 0.5) | 0.000 | 0.380 |
| Waist-to-hip ratio | 0.89 ± 0.06 | 0.91 ± 0.10 | 0.88 ± 0.05 | 0.92 ± 0.10 | 0.00 (-0.02, 0.01) | 0.002 | 0.646 |
| Systolic Blood Pressure (mmHg) | 131 ± 16 | 129 ± 15 | 130 ± 11 | 133 ± 17 | -1.4 (-3.2, 0.4) | 0.009 | 0.123 |
| Diastolic Blood Pressure (mmHg) | 75 ± 10 | 77 ± 10 | 76 ± 10 | 77 ± 10 | -0.5 (-2.0, 1.0) | 0.002 | 0.521 |
| C-reactive protein (mg/L)[b] | 7.6 (9.8) | 4.1 (6.3) | 5.6 (4.4) | 5.4 (4.3) | -1.0 (-1.6, -0.4) | 0.033 | 0.002 |
| White blood cell count (*10^3/mL) | 6.62 ± 1.58 | 6.08 ± 1.73 | 6.27 ± 1.66 | 6.46 ± 1.70 | -0.17 (-0.33, -0.01) | 0.002 | 0.039 |
| Neutrophil-lymphocyte ratio | 2.25 ± 0.65 | 1.89 ± 0.53 | 1.82 ± 0.62 | 2.03 ± 0.67 | -0.12 (-0.20, -0.05) | 0.037 | 0.003 |
| Neutrophil count (*10^3/mL) | 4.03 ± 1.00 | 3.55 ± 1.21 | 3.52 ± 1.14 | 3.88 ± 1.37 | -0.18 (-0.31, -0.05) | 0.022 | 0.008 |
| Lymphocytes count (*10^3/mL) | 1.94 ± 0.74 | 1.92 ± 0.59 | 2.08 ± 0.74 | 1.95 ± 0.49 | 0.01 (-0.06, 0.08) | 0.000 | 0.743 |
| Platelets count (*10^3/mL) | 282 ± 64 | 268 ± 61 | 280 ± 49 | 286 ± 45 | -5 (-10, 0) | 0.008 | 0.053 |
| INFLA score (SD) | 0.08 ± 0.65 | -0.25 ± 0.54 | -0.13 ± 0.50 | -0.03 ± 0.53 | -0.10 (-0.15, -0.04) | 0.030 | <0.001 |
| IL-6 (pg/mL)[b] | 2.06 (0.84) | 2.09 (0.50) | 1.91 (1.02) | 1.80 (1.19) | -0.07 (-0.18, 0.04) | 0.000 | 0.196 |
| Adiponectin (µg/mL) | 5.42 ± 2.79 | 4.70 ± 2.51 | 4.17 ± 2.23 | 4.60 ± 2,60 | -0.29 (-0.53, -0.04) | 0.013 | 0.023 |
| Leptin (ng/ml) | 87.6 ± 37.3 | 90.9 ± 41.1 | 102.0 ± 48.7 | 96.1 ± 45.9 | 1.6 (-1.8, 4.9) | 0.001 | 0.352 |
| HbA1c (mmol/mol) | 37.5 ± 4.0 | 36.8 ± 3.9 | 36.7 ± 4.0 | 36.6 ± 3.8 | -0.2 (-0.4, 0.1) | 0.002 | 0.137 |
| Insulin, fasting (mE/L)[b] | 15.1 (10.1) | 15.0 (13.2) | 18.1 (11.8) | 13.2 (10.2) | 1.1 (0.1, 2.0) | 0.014 | 0.029 |
| Glucose, fasting (mmol/L) | 5.39 ± 0.67 | 5.32 ± 0.40 | 5.48 ± 0.60 | 5.44 ± 0.78 | -0.02 (-0.09, 0.06) | 0.001 | 0.651 |
| HOMA-IR | 3.46 ± 2.02 | 3.83 ± 2.21 | 4.67 ± 2.61 | 3.85 ± 2.20 | 0.24 (-0.01, 0.50) | 0.012 | 0.063 |
| Fatigue (POMS-score, range 0-24)[b] | 6.5 (8.3) | 3.5 (7.0) | 5.0 (5.0) | 3.0 (5.0) | -0.2 (-0.9, 0.5) | 0.002 | 0.578 |
| Depressive mood (POMS-score, range 0-32)[b] | 3.0 (8.0) | 1.5 (4.0) | 1.0 (7.0) | 0.0 (3.0) | -0.2 (-0.6, 0.3) | 0.002 | 0.531 |

*[a]Differences between groups over time were tested by using mixed linear regression, with Time as random factor, and Time, Group, and its interaction as fixed factors of interest. [b]Median (IQR) is reported due to skewed distribution of the variable. BMI, body mass index; HOMA-IR, Homeostatic Model Assessment of Insulin Resistance; IL, interleukin; POMS, Profile of Mood States.*

### Supplementary Table 3: Main task effects of the effort-based decision-making task

| Characteristic | β | 95% CI (β) | OR | 95% CI (OR) | p-value |
|---|---|---|---|---|---|
| EffortLevel | -4.9 | -5.4, -4.4 | 0.007 | 0.004, 0.012 | <0.001 |
| RewardQuantity (high vs low) | 1.9 | 1.6, 2.2 | 6.69 | 4.95, 9.03 | <0.001 |
| Calories (high vs low) | -0.47 | -0.92, -0.02 | 0.63 | 0.40, 0.98 | 0.042 |
| EffortLevel * RewardQuantity | 0.95 | 0.69, 1.2 | 2.59 | 1.99, 3.32 | <0.001 |
| EffortLevel * Calories | 0.23 | 0.02, 0.44 | 1.26 | 1.02, 1.55 | 0.034 |
| RewardQuantity * Calories | -0.13 | -0.25, -0.01 | 0.88 | 0.78, 0.99 | 0.031 |

*Main effects of effort, calories, and reward quantity on offer acceptance, tested by binomial mixed regression modelling (n=148). EffortLevel, RewardQuantity, and Calories and its interactions were added to the model as both fixed and random factors. CI, Confidence Interval; OR, Odds-Ratio.*

### Supplementary Table 4: Whole-brain task effects of the effort-based decision-making task across the cross-sectional and intervention study population

| Contrast | Anatomical region | MNI coordinates | | | *k* | *T-statistic (peak)* | *$P_{FWE}$-value (cluster)* |
|---|---|---|---|---|---|---|---|
| | | x | y | z | | | |
| > Effort | Calcarine, Right | 14 | -92 | 2 | 524 | 9.52 | <0.001 |
| | Occipital Middle Gyrus, Left | -48 | -72 | 0 | 2446 | 8.65 | <0.001 |
| | Superior Medial Frontal Gyrus, Left | 2 | 20 | 42 | 2466 | 7.38 | <0.001 |
| | Insula, Left | -30 | 22 | -10 | 631 | 7.09 | <0.001 |
| | Insula, Right | 38 | 24 | -6 | 468 | 6.46 | <0.001 |
| | Orbitofrontal Middle Frontal Gyrus, Left | -44 | 50 | -10 | 157 | 4.97 | 0.002 |
| | Middle Frontal Gyrus, Right | 44 | 18 | 38 | 294 | 4.87 | <0.001 |
| | Caudate, Left | -14 | 12 | 4 | 140 | 4.77 | 0.005 |
| | Superior Parietal Lobule, Left | -30 | -58 | 58 | 218 | 4.74 | <0.001 |
| | Inferior Parietal Lobule, Right | 54 | -58 | 40 | 316 | 4.60 | <0.001 |
| | Middle Frontal Gyrus, Left | -22 | 52 | 28 | 178 | 4.15 | 0.001 |
| < Effort | Middle Occipital Gyrus, Left | -12 | -94 | 0 | 2464 | 13.03 | <0.001 |
| | Superior Occipital Gyrus, Right | 28 | -80 | 42 | 4693 | 9.68 | <0.001 |
| | Insula, Left | -38 | 0 | 8 | 94 | 5.31 | 0.038 |
| | Middle Frontal Gyrus, Left | -24 | 26 | 44 | 181 | 4.10 | 0.001 |
| High > low calories | Fusiform Gyrus, Right | 32 | -68 | -14 | 2047 | 7.95 | <0.001 |
| | Fusiform Gyrus, Left | -30 | -50 | -12 | 469 | 6.29 | <0.001 |
| | Superior Occipital Gyrus, Left | -16 | -98 | 12 | 178 | 6.02 | 0.001 |
| | Inferior Frontal Gyrus (Triangular Part), Right | 52 | 24 | 6 | 149 | 4.50 | 0.003 |
| Low > high calories | Calcarine, Right | 8 | -90 | 0 | 8418 | 15.31 | <0.001 |
| | Inferior Temporal Gyrus, Left | -48 | -48 | -24 | 88 | 5.04 | 0.05 |
| | Inferior Frontal Gyrus (Triangular Part), Left | -48 | 38 | 16 | 189 | 4.91 | 0.001 |
| High > low reward quantity | Fusiform Gyrus, Right | 26 | -78 | -10 | 8761 | 14.07 | <0.001 |
| | Supramarginal Gyrus, Left | -62 | -54 | 34 | 163 | 4.27 | 0.002 |
| Low > high Reward quantity | Middle Occipital Gyrus, Right | 44 | -76 | 32 | 1082 | 6.39 | <0.001 |
| | Middle Occipital Gyrus, Left | -42 | -70 | 6 | 447 | 5.41 | <0.001 |
| | Lingual Gyrus, Right | 12 | -74 | -2 | 116 | 4.63 | 0.013 |

*Brain regions showing significant effects of effort, reward quantity, and calories at the whole-brain level (FWE-corrected p<0.05 at cluster level). Results are based on one-sample t-tests on the corresponding first-level contrasts.*

### Supplementary Table 5: Task characteristics at baseline

| | **Cross-sectional study (n=131)** | **Intervention study (n=49)** | | |
|---|---|---|---|---|
| | | **Colchicine group (n=24)** | **Placebo group (n=25)** | |
| | ***Mean ± SD or n (%)*** | ***Mean ± SD or n (%)*** | ***Mean ± SD or n (%)*** | ***P-value (colchicine vs placebo)[a]*** |
| Wanting scores low-caloric snack (VAS-score, range 0-1) | 0.60 ± 0.28 | 0.65 ± 0.25 | 0.62 ± 0.27 | 0.691 |
| Wanting scores high-caloric snack (VAS-score, range 0-1) | 0.49 ± 0.29 | 0.43 ± 0.28 | 0.53 ± 0.27 | 0.211 |
| Maximum voluntary contraction (Newton) | 24.2 ± 6.1 | 24.0 ± 5.1 | 24.3 ± 6.5 | 0.858 |
| Success rate of perform trials (%) | 97.2 ± 6.0 | 98.8 ± 3.0 | 97.7 ± 5.3 | 0.395 |
| Earned high-caloric snacks (n of snacks) | 28 ± 9 | 24 ± 10 | 26 ± 11 | 0.457 |
| Earned low-caloric snacks (n of snacks) | 25 ± 12 | 28 ± 12 | 27 ± 10 | 0.774 |

*[a]Difference between intervention groups at baseline was tested by independent t-test for continuous variables and chi-square test for categorical variables. VAS, visual analogue scale*

### Supplementary Table 6: The association between the INFLA score and effort aversion, reward quantity and calorie sensitivity

| Characteristic | β | 95% CI (β) | OR | 95% CI (OR) | p-value |
|---|---|---|---|---|---|
| INFLA | 0.81 | 0.01, 1.6 | 2.25 | 1.01, 4.95 | 0.047 |
| EffortLevel | -4.8 | -5.3, -4.3 | 0.008 | 0.005, 0.014 | <0.001 |
| RewardQuantity (high vs low) | 1.8 | 1.4, 2.1 | 6.05 | 4.05, 8.17 | <0.001 |
| Calories (high vs low) | -0.45 | -0.94, 0.03 | 0.64 | 0.39, 1.03 | 0.068 |
| Age | 0.28 | -0.15, 0.71 | 1.32 | 0.86, 2.03 | 0.200 |
| BMI | 0.02 | -0.43, 0.46 | 1.02 | 0.65, 1.58 | 0.938 |
| INFLA * EffortLevel | -1.3 | -2.1, -0.40 | 0.27 | 0.12, 0.67 | 0.004 |
| INFLA * RewardQuantity | 0.42 | -0.16, 1.0 | 1.52 | 0.85, 2.72 | 0.156 |
| EffortLevel * RewardQuantity | 0.90 | 0.62, 1.2 | 2.46 | 1.86, 3.32 | <0.001 |
| INFLA * Calories | 0.79 | -0.07, 1.6 | 2.20 | 0.93, 4.95 | 0.072 |
| EffortLevel * Calories | 0.22 | -0.01, 0.45 | 1.25 | 0.99, 1.57 | 0.061 |
| RewardQuantity * Calories | -0.15 | -0.27, -0.02 | 0.86 | 0.76, 0.98 | 0.019 |
| INFLA * EffortLevel * RewardQuantity | 0.15 | -0.34, 0.65 | 1.16 | 0.71, 1.91 | 0.546 |
| INFLA * EffortLevel * Calories | -0.06 | -0.50, 0.38 | 0.94 | 0.61, 1.46 | 0.783 |
| INFLA * RewardQuantity * Calories | 0.17 | -0.05, 0.40 | 1.19 | 0.95, 1.49 | 0.136 |

*Associations between the INFLA score and offer acceptance for effort, calories, and reward quantity, tested by binomial mixed regression modelling (n=131). Effort, RewardQuantity, Calories, and its interactions were added as random factors, and INFLA, Effort, Calories, RewardQuantity, and its interactions were added as fixed factors to the model. The model was adjusted by Age and BMI as fixed factors of non-interest. All continuous predictors were Z-scored. CI, Confidence Interval; BMI, body mass index; OR, Odds-Ratio.*

### Supplementary Table 7: The effect of the intervention on effort aversion, reward quantity and calorie sensitivity

| Characteristic | β | 95% CI (β) | OR | 95% CI (OR) | p-value |
|---|---|---|---|---|---|
| Group (colchicine vs placebo) | 0.20 | -0.45, 0.86 | 1.22 | 0.64, 2.36 | 0.542 |
| Timepoint (post vs pre) | -0.01 | -0.38, 0.36 | 0.99 | 0.68, 1.43 | 0.968 |
| EffortLevel | -4.9 | -5.6, -4.3 | 0.007 | 0.004, 0.014 | <0.001 |
| RewardQuantity (high vs low) | 2.4 | 1.9, 3.0 | 11.02 | 6.69, 20.09 | <0.001 |
| Calories (high vs low) | -0.22 | -0.92, 0.47 | 0.80 | 0.40, 1.60 | 0.531 |
| BMI_baseline | 0.44 | -0.21, 1.1 | 1.55 | 0.81, 3.00 | 0.182 |
| Age_baseline | -0.39 | -1.1, 0.27 | 0.68 | 0.33, 1.31 | 0.247 |
| CRP_baseline | -0.39 | -1.1, 0.29 | 0.68 | 0.33, 1.34 | 0.266 |
| Group * Timepoint | 0.04 | -0.33, 0.41 | 1.04 | 0.72, 1.51 | 0.846 |
| Group * EffortLevel | -0.08 | -0.70, 0.55 | 0.92 | 0.50, 1.73 | 0.809 |
| Timepoint * EffortLevel | -0.34 | -0.61, -0.06 | 0.71 | 0.54, 0.94 | 0.018 |
| Group * RewardQuantity | -0.23 | -0.76, 0.31 | 0.79 | 0.47, 1.36 | 0.408 |
| Timepoint * RewardQuantity | 0.29 | 0.05, 0.53 | 1.34 | 1.05, 1.70 | 0.020 |
| EffortLevel * RewardQuantity | 0.64 | 0.47, 0.82 | 1.90 | 1.60, 2.28 | <0.001 |
| Group * Calories | 0.12 | -0.57, 0.82 | 1.13 | 0.57, 2.27 | 0.726 |
| Timepoint * Calories | 0.01 | -0.13, 0.15 | 1.01 | 0.88, 1.16 | 0.888 |
| RewardQuantity * Calories | 0.00 | -0.12, 0.13 | 1.00 | 0.89, 1.14 | 0.964 |
| EffortLevel * Calories | -0.07 | -0.24, 0.11 | 0.93 | 0.79, 1.12 | 0.447 |
| Group * Timepoint * EffortLevel | 0.28 | 0.01, 0.56 | 1.32 | 1.01, 1.75 | 0.044 |
| Group * Timepoint * RewardQuantity | 0.00 | -0.24, 0.24 | 1.00 | 0.79, 1.27 | 0.981 |
| Group* EffortLevel * RewardQuantity | -0.03 | -0.20, 0.15 | 0.97 | 0.82, 1.16 | 0.757 |
| Timepoint * EffortLevel * RewardQuantity | 0.03 | -0.14, 0.20 | 1.03 | 0.87, 1.22 | 0.713 |
| Group * Timepoint * Calories | 0.15 | 0.00, 0.29 | 1.16 | 1.00, 1.34 | 0.047 |
| Group * RewardQuantity * Calories | -0.04 | -0.16, 0.09 | 0.96 | 0.85, 1.10 | 0.580 |
| Timepoint * RewardQuantity * Calories | -0.01 | -0.13, 0.11 | 0.99 | 0.88, 1.12 | 0.835 |
| Group * EffortLevel * Calories | 0.03 | -0.15, 0.20 | 1.03 | 0.86, 1.22 | 0.752 |
| Timepoint * EffortLevel * Calories | 0.00 | -0.16, 0.16 | 1.00 | 0.85, 1.17 | 0.986 |
| Group * Timepoint * EffortLevel * RewardQuantity | 0.11 | -0.05, 0.28 | 1.12 | 0.95, 1.32 | 0.187 |
| Group * Timepoint * RewardQuantity * Calories | -0.08 | -0.20, 0.04 | 0.92 | 0.82, 1.04 | 0.186 |
| Group * Timepoint * EffortLevel * Calories | -0.03 | -0.20, 0.13 | 0.97 | 0.82, 1.14 | 0.674 |

*The effect of colchicine treatment on offer acceptance for effort, calories, and reward quantity over time compared to placebo, tested by mixed binomial regression modelling (n=49). Effort, RewardQuantity, and Calories, Timepoint, and its interactions were added as random factors, and Effort, RewardQuantity, and Calories, Timepoint, Group, and its interactions were added as fixed factors of interest to the model. The model was adjusted by baseline Age, C-reactive protein and BMI as fixed factors of non-interest. All continuous predictors were Z-scored. CI, Confidence Interval; BMI, body mass index; CRP, C-reactive protein; OR, Odd-Ratio.*

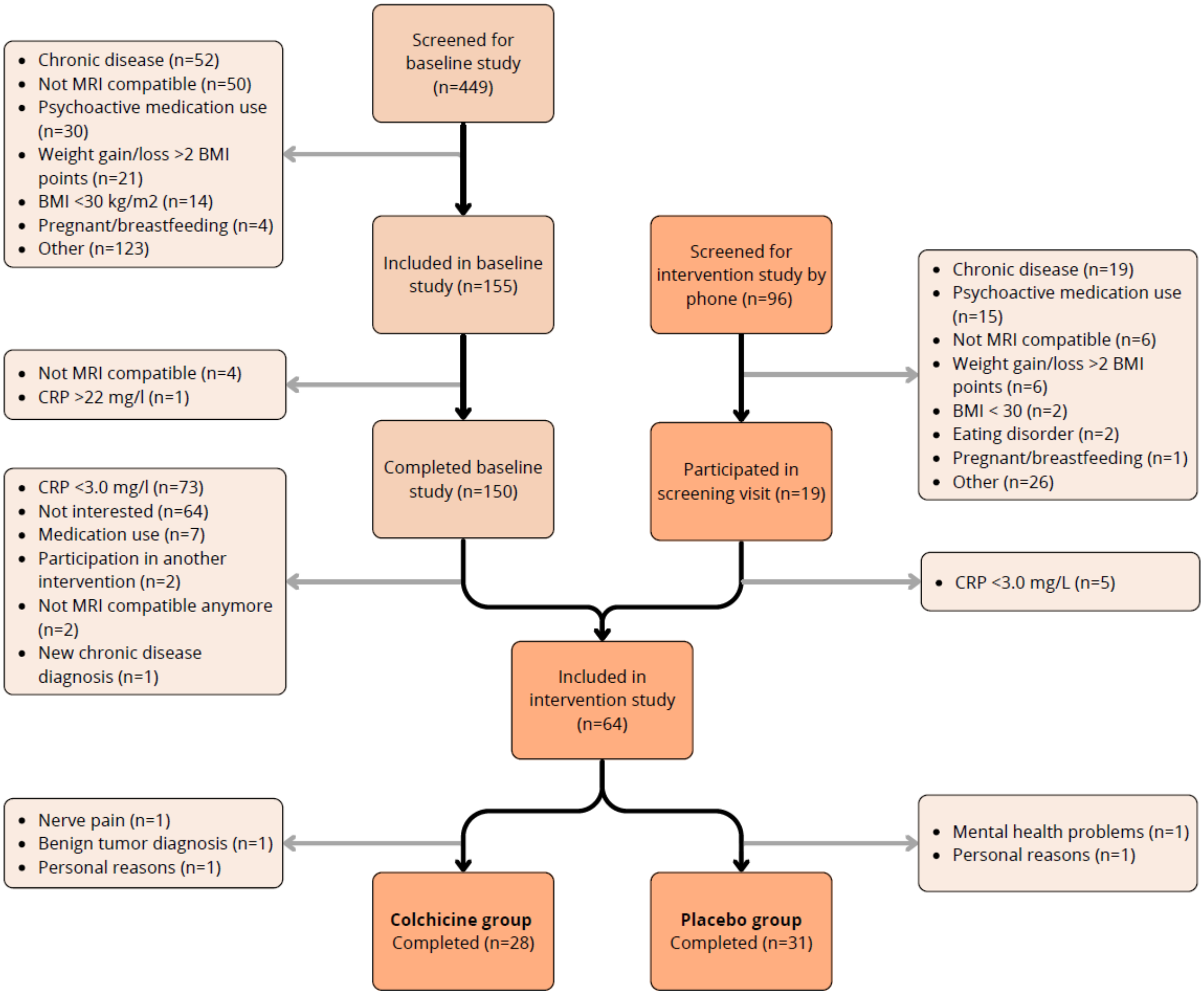

**Supplementary Fig. 1: Flow-chart of the cross-sectional and intervention study population.**

### Supplementary Results: Main behavioural and brain task effects of calorie and reward quantity sensitivity

*Main task effects*

We found main effects for reward quantity (OR=6.69, 95% CI: 4.95 to 9.03, p=<0.001; Fig. S2a) and calories (OR=6.25, 95% CI: 0.40 to 0.98, p=0.042; Fig. S2c), showing that participants accepted more offers with high reward quantity compared to low reward quantity, but unexpectedly less offers with high calories compared to low calories.

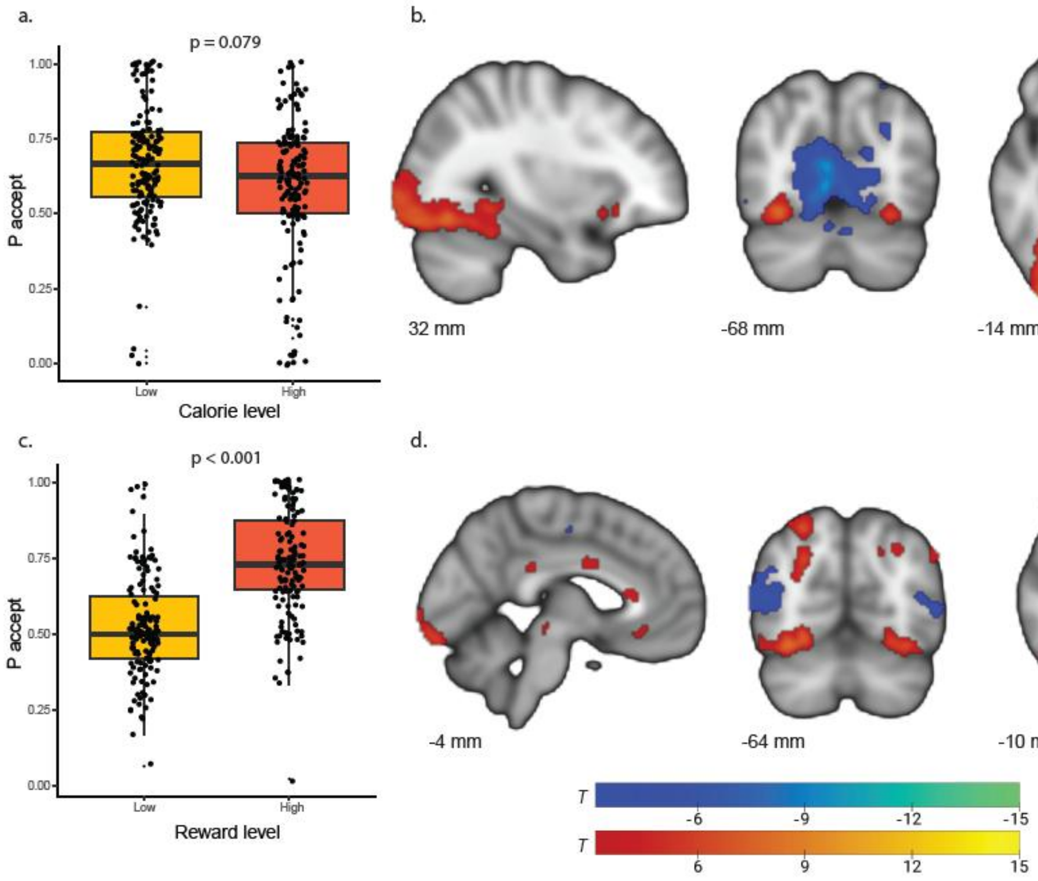

*Fig. S2: Behavioural and whole-brain main task effects for calorie and reward quantity sensitivity (n=148).*

*a, c. Behavioural effects of calories and reward quantity on acceptance rate were tested using mixed binomial regression analysis. b, d. Whole-brain task effects for calories and* *reward quantity are shown at p<0.001 (uncorrected) for illustration purposes. Significant FWE-corrected clusters (p<0.05) are reported in Supplementary material S4.*

*Inflammation and reward quantity and calorie sensitivity*

Using mixed binomial regression analysis, we performed an exploratory analysis to investigate how the INFLA score and colchicine treatment were related to acceptance of offers with high vs. low calories and with high vs. low reward quantity. Higher INFLA score was marginally related to higher calorie sensitivity, i.e. more acceptance of high caloric offers (INFLA*Calories OR=2.20, 95% CI: 0.93 to 4.95, p=0.072; Fig. S3a). The INFLA score was not associated with sensitivity to reward quantity (INFLA*Reward OR=1.52, 95% CI: 0.85 to 2.72, p=0.156; Fig. S3b).

In the intervention study, we found that colchicine increased sensitivity to calories compared with placebo (Group*Timepoint*Calories OR=1.16, 95% CI: 1.00 to 1.34, p=0.047; Fig. S3c). We did not find a difference between groups on reward quantity sensitivity (Group*Timepoint*RewardQuantity OR=1.00 SD, 95% CI: 0.79 to 1.02, p=0.981; Fig. S3d).

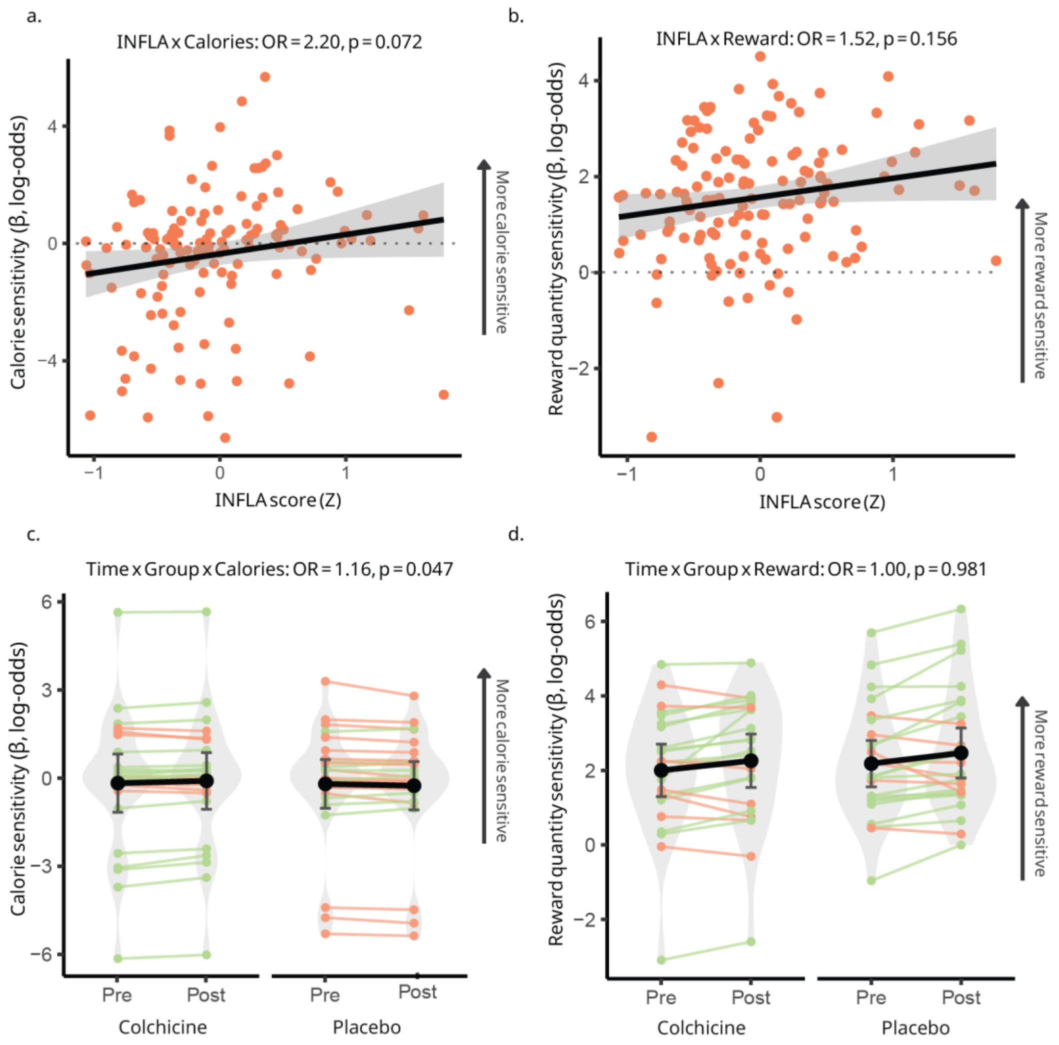

*Fig. S3: The association between the INFLA score and calorie and reward quantity sensitivity, as well as the effect of the intervention on calorie and reward quantity sensitivity.*

*a, b. The relationship between the INFLA score and calorie sensitivity and reward quantity sensitivity, tested by binomial mixed regression modelling. c, d. Average±standard error change (post – pre) in behavioural betas for calorie sensitivity (c) and reward quantity sensitivity (e) according to group, tested by mixed linear regression. OR, odds-ratio.*

### Supplementary Discussion: Inflammation and calorie and reward quantity sensitivity

We explored the role of inflammation on reward sensitivity using two types of reward, caloric content and reward quantity. We did not find any associations between inflammation or colchicine treatment and reward quantity sensitivity. However, regarding calorie sensitivity, we found that colchicine treatment led to an increase in calorie sensitivity. This finding is in line with evidence from mood disorders, where chronic inflammation is related to reward deficits[4,5] and treatment with anti-inflammatory drugs leads to a reduction in reward deficits[6]. While these studies were not able to elucidate effort from reward sensitivity, using an online, non-food version of the same decision-making task as in the current study, we have shown that fatigue complaints in post COVID, likely related to inflammation, are specifically related to reduced reward sensitivity[7]. Together with our findings, this suggests that chronic low-grade inflammation might lead to decreased sensitivity to (food) reward in obesity.

On the other hand, we found that higher inflammation was marginally association with more sensitivity to calories in the cross-sectional study. One possible explanation is a reverse causal relationship, as the cross-sectional design does not allow for causal inferences. One of the factors leading to the development of obesity is thought to be genetically-determined increased sensitivity to food rewards[1,2]. This increased reward sensitivity may promote the intake of high-caloric foods, which in turn can trigger inflammatory responses, particularly when such foods are rich in saturated fats or have a high glycaemic index[3]. Thus, elevated inflammation in our cross-sectional findings might reflect a consequence of increased calorie sensitivity rather than its cause. Taken together, these seemingly contradictory findings highlight the need for longitudinal studies across a broader BMI range to clarify how obesity develops and how sustained inflammation shapes reward and calorie sensitivity.

Supplementary material for: Scholing et al., *Low-grade inflammation in obesity causes low-effort food choice*

### Supplementary Methods 1: In- and exclusion criteria of the cross-sectional and intervention study

#### Cross-sectional study

Inclusion criteria:

- BMI ≥ 27 kg/m2
- Female sex
- Age: 18-59 years
- Shoulder width of < 68 cm (to fit into the MRI scanner)
- Dutch speaking – Sufficient level to understand task instructions
- Willing to comply with the study procedures
- Written informed consent

Exclusion criteria:

- Having been vaccinated by any type of vaccine in the 4 weeks preceding the first test session
- Having had an infection characterized by a fever, or diagnosed by a medical physician in the 4 weeks preceding the first test session, or having CRP levels indicating acute infection (Table S7)
- Diagnosed with Diabetes Mellitus type I or II
- Gained or lost >2 points in BMI (kg/m2) over the last 6 months
- Followed an energy restricting diet during the last 2 months
- Having had bariatric surgery in the past 5 years
- Regular use of anti-inflammatory, anti-diabetic, weight-loss, and psychoactive medication
- (History of) clinically significant psychiatric or neurological disorder
- (History of) clinically significant metabolic, cardiovascular, renal, liver, endocrinological, autoimmune or chronic inflammatory disease
- General medical conditions, such as sensorimotor handicaps, deafness, blindness or colour-blindness, as judged by the investigator
- Current or history of alcohol and/or drugs abuse (i.e. >14 units per week)
- Habitual smoking, i.e. one or more cigarettes per day
- Pregnant, lactating or wishing to become pregnant in the period between the screening and until 3 months after the last study visit (self-reported)
- Allergies or intolerance to any ingredient of the foods used in the study
- Not willing to eat the foods used in this study because of eating habits or believes
- Participation in another weight loss, lifestyle or anti-inflammatory intervention in the context of research at the time of inclusion or during the study
- Contraindications for fMRI:

- IUD as a contraceptive (except MRI-safe IUDs)
- Metal objects or fragments in the body that cannot be taken out
- Active implants in the body
- Using medical plasters
- Epilepsy
- Previous head surgery
- Claustrophobia

*Table S7: Inclusion criteria regarding low-grade inflammation.*

| | **Study** | **First measurement** | **Second measurement at least 4 weeks later** |
|---|---|---|---|
| | | **CRP (mg/L)** | **CRP (mg/L)** |
| **BMI: 30-31 kg/m2** | Cross-sectional | <10.0 | <19.7 |
| | Intervention | 3.0-10.0 | 3.0-19.7 |
| **BMI: >31 kg/m2** | Cross-sectional | <22.1 | <27.8 |
| | Intervention | 3.0-22.1 | 3.0-27.8 |

**Additional criteria for the intervention study:**

Inclusion criteria

- BMI ≥ 30 kg/m2
- Low-grade inflammatory state, as measured by CRP according to BMI and type of measurement. Table S7 below shows an overview of the CRP inclusion criteria.

Exclusion criteria

- Regular use of CYP3A4 inhibitors, P-glycoprotein inhibitors, statins, fibrates, ciclosporin, and digoxin, as a contraindication for colchicine. A list of the most commonly used medication that interacts with colchicine can be found in Appendix A.
- Having renal impairment as evidenced by serum creatinine >150 µmol/l or eGFR <50mL/min/1.73m2, determined maximum 12 weeks before inclusion.
- Having moderate to severe hepatic disease

### Supplementary Methods 2: Description of the study procedure

*Procedure*

After we recruited participants for the cross-sectional study, we invited them for a 4.5h visit at the study centre. We asked participants to refrain from eating and drinking anything other than water 12 hours before the test session. All visits were planned in the morning with a start time between 8-11:30am. Upon arrival at the centre, participants signed informed consent and we measured CRP via a finger prick to screen for acute infections. In addition, the weight of the participant was measured using a calibrated scale. If all inclusion criteria were met, we drew a venous blood sample, measured the participants' blood pressure, and waist and hip circumference, and took a hair sample for the measurement of cortisol. In addition, participants were asked to collect their second morning urine sample. Hereafter, we gave the participant a yoghurt drink as standardized breakfast. Then, participants performed cognitive tests (digit span and Stroop task) and filled in a range of questionnaires assessing their lifestyle, moods and behaviours (see Table S8). Hereafter, participants underwent 1.5h of MRI scanning, which included anatomical imaging, magnetic resonance spectroscopy (MRS), and three sets of functional MRI brain sequences: a food-related effort-based decision-making task, a resting state scan and a monetary incentive delay task. In addition, we acquired an abdominal anatomical scan to measure subcutaneous and visceral fat. After scanning, participants performed a food-intake test to measure effort- and reward-related food intake.

In the week after the visit, participants collected a faecal sample at home for the analysis of microbiome composition and metabolite profiling. In addition, participants completed an experience sampling method (ESM)-protocol, for which they filled out 8 brief questionnaires per day on their mobile phone for a period of 8-10 days to assess motivational behaviour in daily life. That concluded their participation in the cross-sectional study.

Participants that were included in the intervention study were invited for a screening visit at the test centre, during which participants signed informed consent and we took a capillary blood sample by a finger prick to check whether participants met the criteria regarding low-grade inflammation and kidney function. If all inclusion criteria were met, participants collected a faecal sample at home, and performed the same ESM measurement as described above. Hereafter, the participant attended a baseline study visit at the test centre, which was identical to the visit described above.

For participants who first took part in the cross-sectional study and later enrolled in the intervention study, eligibility for the intervention was assessed during their test session in the cross-sectional study. For these participants, this test session and the home measurements also served as the baseline measurements for the intervention study.

After completion of the baseline visit and home measurement, participants were randomized into the colchicine or placebo group and received the study medication. During the intervention period of 12 weeks, we called the participants three time (in week 2, 4 and 8) to check for potential adverse events and compliance to the intervention. In the 11$^{th}$ week of the intervention, participants collected a faecal sample again at home, and did an ESM measurement for 8-10 days again. In the 12th and final week of the intervention period, participants attended a follow-up visit at the test centre with the same starting time as their baseline visit. The follow-up visit was identical to the baseline visit.

*Table S8: Questionnaires included in the study*

| **Name** | **Baseline** | **Follow-up** |
|---|---|---|
| Dutch Eating Behaviour Questionnaire (DEBQ)[8] | X | |
| Binge Eating Scale (BES)[9] | X | X |
| Power of Food Scale (PoFS)[10] | X | X |
| Food Frequency Questionnaire (FFQ)[11] | X | X |
| Hospital Anxiety and Depression Scale (HADS)[12] | X | X |
| Checklist Individual Strength (CIS)[13] | X | X |
| Profile of Mood State (POMS)[14] | X | X |
| 36-Item Short Form Survey (SF-36)[15] | X | X |
| Temporal Experience of Pleasure Scale (TEPS)[16] | X | X |
| Starkstein Apathy Scale (SAS)[17] | X | X |

| | | |
|---|---|---|
| Barratt Impulsivity Scale[18] | X | |
| Behavioural Activation System/ Behavioural Inhibition System (BIS/BAS) Scale[19] | X | X |
| Kirby's Monetary Choice Questionnaire[20,21] | X | X |
| Pittsburgh Sleep Quality Index (PSQI)[22] | X | X |
| Demographic and lifestyle questionnaire | X | |
| Menstrual cycle questionnaire | X | X |
| Fasting questionnaire | X | X |

### Supplementary Methods 3: Functional MRI pre-processing pipeline using fmriprep.

*Anatomical data preprocessing*

A total of 2 T1-weighted (T1w) images were found within the input BIDS dataset. Each T1w image was corrected for intensity non-uniformity (INU) with N4BiasFieldCorrection (Tustison et al. 2010), distributed with ANTs 2.5.1 (Avants et al. 2008, RRID:SCR_004757). The T1w-reference was then skull-stripped with a *Nipype* implementation of the antsBrainExtraction.sh workflow (from ANTs), using OASIS30ANTs as target template. Brain tissue segmentation of cerebrospinal fluid (CSF), white-matter (WM) and gray-matter (GM) was performed on the brain-extracted T1w using fast (FSL (version unknown), RRID:SCR_002823, Zhang, Brady, and Smith 2001). An anatomical T1w-reference map was computed after registration of 2 T1w images (after INU-correction) using mri_robust_template (FreeSurfer 7.3.2, Reuter, Rosas, and Fischl 2010). Brain surfaces were reconstructed using recon-all (FreeSurfer 7.3.2, RRID:SCR_001847, Dale, Fischl, and Sereno 1999), and the brain mask estimated previously was refined with a custom variation of the method to reconcile ANTs-derived and FreeSurfer-derived segmentations of the cortical gray-matter of Mindboggle (RRID:SCR_002438, Klein et al. 2017). Volume-based spatial normalization to two standard spaces (MNI152NLin6Asym, MNI152NLin2009cAsym) was performed through nonlinear registration with antsRegistration (ANTs 2.5.1), using brain-extracted versions of both T1w reference and the T1w template. The following templates were selected for spatial normalization and accessed with *TemplateFlow* (24.2.0, Ciric et al. 2022): *FSL's MNI ICBM 152 non-linear 6th Generation Asymmetric Average Brain Stereotaxic Registration Model* [Evans et al. (2012), RRID:SCR_002823; TemplateFlow ID: MNI152NLin6Asym], *ICBM 152 Nonlinear Asymmetrical template version 2009c* [Fonov et al. (2009), RRID:SCR_008796; TemplateFlow ID: MNI152NLin2009cAsym].

*Functional data preprocessing*

For each of the 6 BOLD runs found per subject (across all tasks and sessions), the following preprocessing was performed. First, a reference volume was generated, using a custom methodology of *fMRIPrep*, for use in head motion correction. Head-motion parameters with respect to the BOLD reference (transformation matrices, and six corresponding rotation and translation parameters) are estimated before any spatiotemporal filtering using mcflirt (FSL , Jenkinson et al. 2002). The BOLD reference was then co-registered to the T1w reference using bbregister (FreeSurfer) which implements boundary-based registration (Greve and Fischl 2009). Co-registration was conFig.d with six degrees of freedom. Several confounding time-series were calculated based on the *preprocessed BOLD*: framewise displacement (FD), DVARS and three region-wise global signals. FD was computed using two formulations following Power (absolute sum of relative motions, Power et al. (2014)) and Jenkinson (relative root mean square displacement between affines, Jenkinson et al. (2002)). FD and DVARS are calculated for each functional run, both using their implementations in *Nipype* (following the definitions by Power et al. 2014). The three global signals are extracted within the CSF, the WM, and the whole-brain masks. Additionally, a set of physiological regressors were extracted to allow for component-based noise correction (*CompCor*, Behzadi et al. 2007). Principal components are estimated after high-pass filtering the *preprocessed BOLD* time-series (using a discrete cosine filter with 128s cut-off) for the two *CompCor* variants: temporal (tCompCor) and anatomical (aCompCor). tCompCor components are then calculated from the top 2% variable voxels within the brain mask. For aCompCor, three probabilistic masks (CSF, WM and combined CSF+WM) are generated in anatomical space. The implementation differs from that of Behzadi et al. in that instead of eroding the masks by 2 pixels on BOLD space, a mask of pixels that likely contain a volume fraction of GM is subtracted from the aCompCor masks. This mask is obtained by dilating a GM mask extracted from the FreeSurfer's *aseg* segmentation, and it ensures components are not extracted from voxels containing a minimal fraction of GM. Finally, these masks are resampled into BOLD space and binarized by thresholding at 0.99 (as in the original implementation). Components are also calculated separately within the WM and CSF masks. For each CompCor decomposition, the *k* components with the largest singular values are retained, such that the retained components' time series are sufficient to explain 50 percent of variance across the nuisance mask (CSF, WM, combined, or temporal). The remaining components are dropped from consideration. The head-motion estimates calculated in the correction step were also placed within the corresponding confounds file. The confound time series derived from head motion estimates and global signals were expanded with the inclusion of temporal derivatives and quadratic terms for each (Satterthwaite et al. 2013). Frames that exceeded a threshold of 0.5 mm FD or 1.5 standardized DVARS were annotated as motion outliers. Additional nuisance timeseries are calculated by means of principal components analysis of the signal found within a thin band (*crown*) of voxels around the edge of the brain, as proposed by (Patriat, Reynolds, and Birn 2017). All resamplings can be performed with *a single interpolation step* by composing all the pertinent transformations (i.e. head-motion transform matrices, susceptibility distortion correction when available, and co-registrations to anatomical and output spaces). Gridded (volumetric) resamplings were performed using nitransforms, conFig.d with cubic B-spline interpolation.

Many internal operations of *fMRIPrep* use *Nilearn* 0.10.4 (Abraham et al. 2014, RRID:SCR_001362), mostly within the functional processing workflow. For more details of the pipeline, see the section corresponding to workflows in *fMRIPrep*'s documentation.

*Copyright Waiver*

The above boilerplate text was automatically generated by fMRIPrep with the express intention that users should copy and paste this text into their manuscripts *unchanged*. It is released under the CC0 license.

*References*

Abraham, Alexandre, Fabian Pedregosa, Michael Eickenberg, Philippe Gervais, Andreas Mueller, Jean Kossaifi, Alexandre Gramfort, Bertrand Thirion, and Gael Varoquaux.

2014. "Machine Learning for Neuroimaging with Scikit-Learn." *Frontiers in Neuroinformatics* 8. https://doi.org/10.3389/fninf.2014.00014.

Avants, B. B., C. L. Epstein, M. Grossman, and J. C. Gee. 2008. "Symmetric Diffeomorphic Image Registration with Cross-Correlation: Evaluating Automated Labeling of Elderly and Neurodegenerative Brain." *Medical Image Analysis* 12 (1): 26–41. https://doi.org/10.1016/j.media.2007.06.004.

Behzadi, Yashar, Khaled Restom, Joy Liau, and Thomas T. Liu. 2007. "A Component Based Noise Correction Method (CompCor) for BOLD and Perfusion Based fMRI." *NeuroImage* 37 (1): 90–101. https://doi.org/10.1016/j.neuroimage.2007.04.042.

Ciric, R., William H. Thompson, R. Lorenz, M. Goncalves, E. MacNicol, C. J. Markiewicz, Y. O. Halchenko, et al. 2022. "TemplateFlow: FAIR-Sharing of Multi-Scale, Multi-Species Brain Models." *Nature Methods* 19: 1568–71. https://doi.org/10.1038/s41592-022-01681-2.

Dale, Anders M., Bruce Fischl, and Martin I. Sereno. 1999. "Cortical Surface-Based Analysis: I. Segmentation and Surface Reconstruction." *NeuroImage* 9 (2): 179–94. https://doi.org/10.1006/nimg.1998.0395.

Esteban, Oscar, Ross Blair, Christopher J. Markiewicz, Shoshana L. Berleant, Craig Moodie, Feilong Ma, Ayse Ilkay Isik, et al. 2018. "fMRIPrep 24.0.0." *Software*. https://doi.org/10.5281/zenodo.852659.

Esteban, Oscar, Christopher Markiewicz, Ross W Blair, Craig Moodie, Ayse Ilkay Isik, Asier Erramuzpe Aliaga, James Kent, et al. 2019. "fMRIPrep: A Robust Preprocessing Pipeline for Functional MRI." *Nature Methods* 16: 111–16. https://doi.org/10.1038/s41592-018-0235-4.

Evans, AC, AL Janke, DL Collins, and S Baillet. 2012. "Brain Templates and Atlases." *NeuroImage* 62 (2): 911–22. https://doi.org/10.1016/j.neuroimage.2012.01.024.

Fonov, VS, AC Evans, RC McKinstry, CR Almli, and DL Collins. 2009. "Unbiased Nonlinear Average Age-Appropriate Brain Templates from Birth to Adulthood." *NeuroImage* 47, Supplement 1: S102. https://doi.org/10.1016/S1053-8119(09)70884-5.

Gorgolewski, K., C. D. Burns, C. Madison, D. Clark, Y. O. Halchenko, M. L. Waskom, and S. Ghosh. 2011. "Nipype: A Flexible, Lightweight and Extensible Neuroimaging Data Processing Framework in Python." *Frontiers in Neuroinformatics* 5: 13. https://doi.org/10.3389/fninf.2011.00013.

Gorgolewski, Krzysztof J., Oscar Esteban, Christopher J. Markiewicz, Erik Ziegler, David Gage Ellis, Michael Philipp Notter, Dorota Jarecka, et al. 2018. "Nipype." *Software*. https://doi.org/10.5281/zenodo.596855.

Greve, Douglas N, and Bruce Fischl. 2009. "Accurate and Robust Brain Image Alignment Using Boundary-Based Registration." *NeuroImage* 48 (1): 63–72. https://doi.org/10.1016/j.neuroimage.2009.06.060.

Jenkinson, Mark, Peter Bannister, Michael Brady, and Stephen Smith. 2002. "Improved Optimization for the Robust and Accurate Linear Registration and Motion Correction of Brain Images." *NeuroImage* 17 (2): 825–41. https://doi.org/10.1006/nimg.2002.1132.

Klein, Arno, Satrajit S. Ghosh, Forrest S. Bao, Joachim Giard, Yrjö Häme, Eliezer Stavsky, Noah Lee, et al. 2017. "Mindboggling Morphometry of Human Brains." *PLOS Computational Biology* 13 (2): e1005350. https://doi.org/10.1371/journal.pcbi.1005350.

Patriat, Rémi, Richard C. Reynolds, and Rasmus M. Birn. 2017. "An Improved Model of Motion-Related Signal Changes in fMRI." *NeuroImage* 144, Part A (January): 74–82. https://doi.org/10.1016/j.neuroimage.2016.08.051.

Power, Jonathan D., Anish Mitra, Timothy O. Laumann, Abraham Z. Snyder, Bradley L. Schlaggar, and Steven E. Petersen. 2014. "Methods to Detect, Characterize, and Remove Motion Artifact in Resting State fMRI." *NeuroImage* 84 (Supplement C): 320–41. https://doi.org/10.1016/j.neuroimage.2013.08.048.

Reuter, Martin, Herminia Diana Rosas, and Bruce Fischl. 2010. "Highly Accurate Inverse Consistent Registration: A Robust Approach." *NeuroImage* 53 (4): 1181–96. https://doi.org/10.1016/j.neuroimage.2010.07.020.

Satterthwaite, Theodore D., Mark A. Elliott, Raphael T. Gerraty, Kosha Ruparel, James Loughead, Monica E. Calkins, Simon B. Eickhoff, et al. 2013. “An improved framework for confound regression and filtering for control of motion artifact in the preprocessing of resting-state functional connectivity data.” *NeuroImage* 64 (1): 240–56. https://doi.org/10.1016/j.neuroimage.2012.08.052.

Tustison, N. J., B. B. Avants, P. A. Cook, Y. Zheng, A. Egan, P. A. Yushkevich, and J. C. Gee. 2010. “N4ITK: Improved N3 Bias Correction.” *IEEE Transactions on Medical Imaging* 29 (6): 1310–20. https://doi.org/10.1109/TMI.2010.2046908.

Zhang, Y., M. Brady, and S. Smith. 2001. “Segmentation of Brain MR Images Through a Hidden Markov Random Field Model and the Expectation-Maximization Algorithm.” *IEEE Transactions on Medical Imaging* 20 (1): 45–57. https://doi.org/10.1109/42.906424.

### References

1. Farooqi, S. Obesity and thinness: insights from genetics. *Philos Trans R Soc Lond B Biol Sci* **378**, 20220205 (2023).
2. Rapuano, K. M. *et al.* Genetic risk for obesity predicts nucleus accumbens size and responsivity to real-world food cues. *Proc Natl Acad Sci U S A* **114**, 160–165 (2017).
3. Malesza, I. J. *et al.* High-Fat, Western-Style Diet, Systemic Inflammation, and Gut Microbiota: A Narrative Review. *Cells* **10**, 3164 (2021).
4. Capuron, L. & Miller, A. H. Cytokines and psychopathology: lessons from interferon-alpha. *Biol Psychiatry* **56**, 819–824 (2004).
5. Felger, J. C. Imaging the Role of Inflammation in Mood and Anxiety-related Disorders. *Curr Neuropharmacol* **16**, 533–558 (2018).
6. Raison, C. L. *et al.* A randomized controlled trial of the tumor necrosis factor antagonist infliximab for treatment-resistant depression: the role of baseline inflammatory biomarkers. *JAMA Psychiatry* **70**, 31–41 (2013).
7. Scholing, J. M., Lambregts, B. I. H. M., van den Bosch, R., Aarts, E. & van der Schaaf, M. E. Greater fatigue is more strongly associated with reduced reward sensitivity in the long-term phase of coronavirus disease (COVID-19) than in the early phase. *Brain Behav Immun Health* **48**, 101056 (2025).
8. Cebolla, A., Barrada, J. R., van Strien, T., Oliver, E. & Baños, R. Validation of the Dutch Eating Behavior Questionnaire (DEBQ) in a sample of Spanish women. *Appetite* **73**, 58–64 (2014).

9. The assessment of binge eating severity among obese persons - PubMed. https://pubmed.ncbi.nlm.nih.gov/7080884/.

10. Lowe, M. R. *et al.* The Power of Food Scale. A new measure of the psychological influence of the food environment. *Appetite* **53**, 114–118 (2009).

11. van Lee, L. *et al.* The Dutch Healthy Diet index as assessed by 24 h recalls and FFQ: associations with biomarkers from a cross-sectional study. *J Nutr Sci* **2**, e40 (2013).

12. Zigmond, A. S. & Snaith, R. P. The hospital anxiety and depression scale. *Acta Psychiatr Scand* **67**, 361–370 (1983).

13. The assessment of fatigue: Psychometric qualities and norms for the Checklist individual strength - PubMed. https://pubmed.ncbi.nlm.nih.gov/28554371/.

14. McNair et al. *Manual for the Profile of Mood States.* (Educational and Industrial Testing Service., San Diego, CA, 1971).

15. Ware, J. E. & Sherbourne, C. D. The MOS 36-item short-form health survey (SF-36). I. Conceptual framework and item selection. *Med Care* **30**, 473–483 (1992).

16. Simon, J. J. *et al.* Psychometric evaluation of the Temporal Experience of Pleasure Scale (TEPS) in a German sample. *Psychiatry Res* **260**, 138–143 (2018).

17. Starkstein, S. E. *et al.* Reliability, validity, and clinical correlates of apathy in Parkinson's disease. *J Neuropsychiatry Clin Neurosci* **4**, 134–139 (1992).

18. Patton, J. H., Stanford, M. S. & Barratt, E. S. Factor structure of the Barratt impulsiveness scale. *J Clin Psychol* **51**, 768–774 (1995).

19. Carver, C. S. & White, T. L. Behavioral inhibition, behavioral activation, and affective responses to impending reward and punishment: The BIS/BAS Scales. *Journal of Personality and Social Psychology* **67**, 319–333 (1994).

20. Kirby, K. N., Petry, N. M. & Bickel, W. K. Heroin addicts have higher discount rates for delayed rewards than non-drug-using controls. *J Exp Psychol Gen* **128**, 78–87 (1999).

21. Kirby, K. N. & Maraković, N. N. Delay-discounting probabilistic rewards: Rates decrease as amounts increase. *Psychon Bull Rev* **3**, 100–104 (1996).

22. Buysse, D. J., Reynolds, C. F., Monk, T. H., Berman, S. R. & Kupfer, D. J. The Pittsburgh Sleep Quality Index: a new instrument for psychiatric practice and research. *Psychiatry Res* **28**, 193–213 (1989).

23. Tustison, N. J. *et al.* N4ITK: Improved N3 Bias Correction. *IEEE Transactions on Medical Imaging* **29**, 1310–1320 (2010).
